# Adaptive gene transcription in *Escherichia coli* under environmental stress

**DOI:** 10.64898/2026.02.10.705075

**Authors:** Pingzhuang Ge, Fatema-Zahra M. Rashid, Luuk K. F. Gaarthuis, Marc K. M. Cajili, Minkang Tan, Baoxu Pang, Karin Schnetz, Remus T. Dame

## Abstract

*Escherichia coli* is highly sensitive to acid and osmotic stress but adapts by modulating the expression of stress responsive genes. Nucleoid-associated proteins (NAPs) play key roles in DNA organization and sensing environmental changes. The histone-like nucleoid structuring protein H-NS is an NAP acting as a global regulator of stress genes. H-NS may alter local chromatin structure to modulate the expression of such genes in response to environmental stress. The H-NS homolog StpA co-regulates several target genes, but its precise role is poorly defined. To investigate the regulatory interplay between these two proteins, we examined transcription, DNA binding and chromatin structure at two regulated operons, *hdeAB* and *proVWX, in E. coli* following exposure to acid and salt shock. Our results show that H-NS senses pH and osmotic cues to remodel chromatin and relieve repression, while StpA compensates for H-NS loss, particularly at *proVWX*, highlighting a coordinated regulatory network.

## Introduction

Bacteria inhabit complex and changing environments, resulting in physico-chemical stress arising from changes in pH, temperature, osmotic pressure and nutrient availability. Large environmental changes can pose serious threats to bacterial survival. Under stressful conditions, bacteria rapidly sense and adapt to changes in the extracellular environment to maintain intracellular homeostasis and ensure survival. *Escherichia coli* serves as a well-studied example, with extensive research demonstrating that it has evolved powerful regulatory systems to respond to a wide range of environmental stresses (1–5). However, how *E. coli* selectively responds to environmental stresses to regulate only the specific subset of genes required for that stress response is still largely unclear.

Nucleoid associated proteins (NAPs) shape chromosome architecture and regulate global transcription in bacteria and thus play a crucial role in response to environmental change (6). The histone-like nucleoid structuring protein (H-NS) (7–9) is a small NAP widespread in gram-negative bacteria (10–12). H-NS consists of three functional domains: a C-terminal DNA binding domain, an N-terminal dimerization domain and an internal dimer-dimer interaction domain (12, 13). H-NS multimerizes along DNA to form a lateral H-NS-DNA filament or, when recruiting a second DNA segment, a bridged H-NS-DNA complex (6, 12, 14–17). Binding in trans, DNA bridging depends on availability of the two DNA binding domains in the H-NS dimer, which is subject to regulation (14, 18). H-NS exhibits preferential binding to AT-rich DNA regions (19–23). Occupancy of promoter regions by H-NS hinders the binding of RNA polymerase (24–27). The formation of intragenic H-NS–DNA bridges impedes RNA polymerase movement along the DNA (18, 26, 28). In both cases this induces repression of transcription in either the transcription initiation or transcription elongation phase (9). In *E. coli*, around 5% of the genes are silenced by H-NS directly. This includes many genes and operons involved in environmental response, for example, the acid resistance operon *hdeAB* and the osmotic resistance operon *proVWX* (29–33). Often, genomic regions targeted by H-NS have been acquired via horizontal gene transfer (34–37). Over the past two decades, numerous anti-silencing mechanisms involving antagonistic proteins have been uncovered (38–49). Additionally, many studies have shown that the protein structure, multimerization, post-translational modifications (PTMs) and DNA binding ability of H-NS can be modulated by external environmental stimuli (6, 14, 22, 37, 50–52). Together, these observations indicate that H-NS is a direct sensor of physico-chemical signals. As an enterobacterium, *E. coli* inevitably faces the challenge of acid stress in its host’s digestive system (for example, the pH is 2.0 in the human stomach). To cope with this challenge, a dedicated acid fitness island (AFI) containing 13 acid resistance genes is harbored in the *E. coli* genome (53, 54). Genome-wide ChIP data have shown that H-NS exhibits widespread binding across this genomic island (19–23). Such binding of H-NS underlies the role of H-NS in repression of these acid resistance genes (29). The mechanism of repression by H-NS and the relief of repression following acid shock is largely unclear.

StpA (suppressor of td phenotype A), a homolog of H-NS, shares 58% sequence identity with H-NS and exhibits a highly similar overall protein structure (55–58). *E. coli* contains around 47, 000 copies of H-NS and 5, 200 copies of StpA per cell in exponential phase (59, 60). *In vitro* studies have shown that StpA can form heterodimers with H-NS to create a DNA–H-NS/StpA bridging complex (61–64) and can also independently form filaments or DNA-bridging structures (65). In contrast to the H-NS–DNA complex, the StpA–DNA complex exhibits minimal responsiveness to alterations in salt concentration, temperature and pH *in vitro* (65). Moreover, StpA and H-NS negatively regulate each other’s transcription (57, 66). The current consensus is that StpA is a molecular back-up of H-NS, which takes over the transcription repression of StpA/H-NS dual target genes (such as, *papB*, *proVWX* and *bgl*) in a Δ*hns* strain (57). However, the molecular mechanism by which StpA responds to environmental stimuli in *E. coli* and cooperates with H-NS to regulate shared target gene transcription remains largely unexplored.

The *hdeAB* operon encodes two periplasmic chaperone proteins: HdeA (67, 68) and HdeB (69). At neutral pH conditions, both HdeA and HdeB exist as inactive homodimers (70). Upon acid stress, they undergo monomerization and conformational changes that expose their hydrophobic surfaces, enabling them to interact with substrate proteins (68, 70). This interaction prevents the denaturation and aggregation of substrate proteins, which is essential for *E. coli* survival under acidic conditions (70). The transcriptional regulatory network of *hdeAB* is highly complex and is modulated by multiple regulators, including RpoD (σ⁷⁰) (71), RpoS (σ³⁸) (71–76), H-NS (23, 33, 71, 76, 77), StpA (23), FliZ (78), ppGpp (79), GadE (53, 75), GadX (75, 77, 80, 81), GadW (75, 80, 81), GadY (82), EvgAS (83), YdeO (83), Lrp (84) and MarA (84, 85). The transcription of *hdeAB* is further influenced by growth phase (71, 73) and the pH of the environment (29, 33, 74). Current evidence suggests that H-NS may occupy the top level of this transcriptional coordination hierarchy (84).

The *proVWX* operon encodes three proteins: ProV, ProW and ProX, which assemble into the ProU transporter (86, 87). Upon exposure to hyperosmotic stress, this transporter mediates the uptake of compatible solutes such as glycine betaine, proline betaine and proline (87–91). These osmoprotectants maintain a low cytoplasmic osmotic potential without exerting detrimental effects on cellular physiology (92, 93). In our previous work, we provided a detailed characterization of the transcriptional regulatory network governing the *proVWX* operon (18). It is particularly noteworthy that the operon is repressed by H-NS and is subject to transcription activation under osmotic stress (18, 23, 88, 94, 95). Recent studies have revealed the mechanism by which H-NS repression of *proVWX* is relieved: under osmotic stress conditions, H-NS undergoes conformational changes at the *proVWX* promoter region, thereby releasing its bridging-mediated transcriptional repression through the reorganization of local three-dimensional chromatin architecture (14, 18).

To investigate the transcriptional regulatory roles of H-NS and StpA in the response of *E. coli* to environmental stress, we subjected the cells to acid or salt shock treatments. RT-qPCR was employed to profile the transcriptional dynamics of the *hdeAB* and *proVWX* operons under varying pH and osmotic conditions. ChIP-qPCR was then used to map the binding patterns of H-NS and StpA at the promoter regions of these target operons, while 3C-qPCR analyses were used to elucidate the local three-dimensional chromatin architecture. Together, these results enabled us to develop models describing the cooperative roles of H-NS and StpA in transcriptional control during the adaptive response of *E. coli* to environmental stimuli.

## Material and methods

### Strain construction

All the *E. coli* strains used in this study were derived from *Escherichia coli* MG1655. All the mutation strains we construct by using two-step genome editing method based on the λ-red recombinase system (96) and phage P1 transduction method (97). The details of construction of strains by using λ-red recombinase system was described in our previous study (18). Strains used in the current study are listed in Table 1.

**Table 1:**
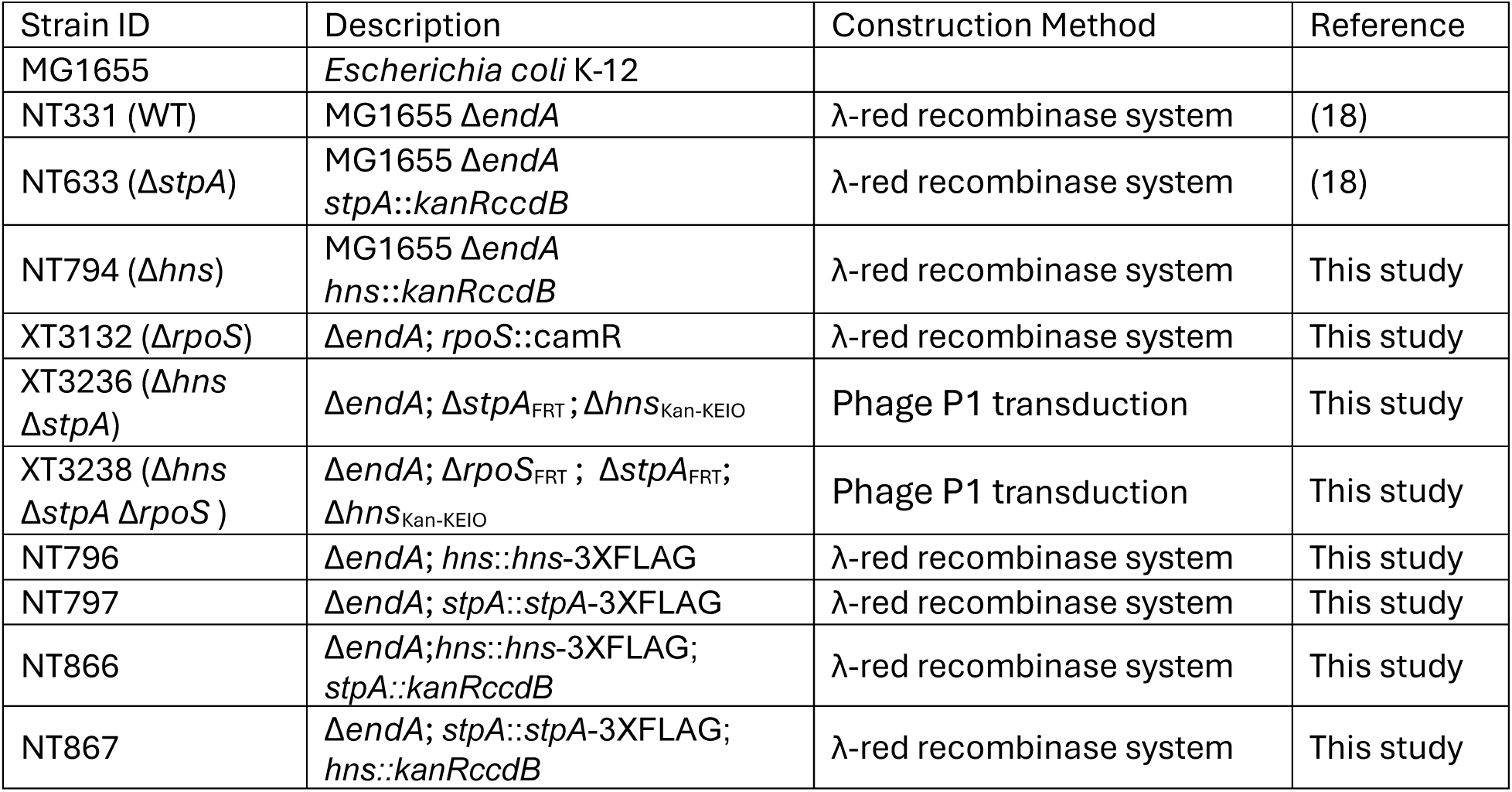
Strains used in this study.

### Growth curves

Individual colonies of *E. coli* strains were cultured overnight in M9 glycerol medium at 37 °C with 200 rpm shaking incubator (Eppendorf Innova® 40) to the stationary phase and then diluted with fresh M9 glycerol medium to an OD_600_ of approximately 0.05. 300 μL of cultures were transferred into 100-well Honeycomb (Catalog number: 95025BIO), each culture with 3 technical replicates. Incubated at 37 °C with continuous shaking using the Bioscreen G Pro™ automated growth analyzer. OD_600_ values were recorded automatically every 30 mins, and growth curves were plotted based on the collected data (**Supplementary figure 1**).

### Growth conditions

All the strains were cultured overnight on LB agar plates with appropriate antibiotic at 37 °C in an incubator (Catalog numbers: 390-0384, VWR® INCU-Line®).

For acid resistance studies, four single colonies were taken from the agar plate and were separately grown overnight in 2 mL M9 glycerol medium (pH=6.6, 42 mM Na_2_HPO_4_, 22 mM KH_2_PO_4_, 19 mM NH_4_Cl, 2.0 mM MgSO_4_, 0.1 mM CaCl_2_, 170 mM NaCl, 1X trace elements, 1% Bacto™ casamino acids (BD), 10 μg/mL thiamine (Sigma-Aldrich), 0.4% glycerol (PanReac Applichem) with proper antibiotic at 37 °C with 200 rpm shaking incubator (Eppendorf Innova® 40). The overnight culture was used to inoculate M9 glycerol medium to a starting OD₆₀₀ of 0.05. The culture was grown in an incubator at 37°C with shaking at 200 rpm until an OD₆₀₀ of 1.0 (early exponential phase). For testing at pH 6.6, 1 mL of the suspension was kept aside for RNA isolation, 9 mL was used for 3C-based studies, 10 mL used for ChIP-based study. For pH 2.0, 37% HCl was added to adjust the pH of the M9 medium to 2.0. The bacterial suspension was then incubated at 37°C with shaking at 200 rpm for 10 minutes.

For osmolarity experiments, four single colonies were taken from the agar plate and were separately grown overnight in 2 mL low salt (LS) M9 glycerol medium (42 mM Na_2_HPO_4_, 22 mM KH_2_PO_4_, 19 mM NH_4_Cl, 2.0 mM MgSO_4_, 0.1 mM CaCl_2_, 80 mM NaCl, 1X trace elements, 1% Bacto™ casamino acids (BD), 10 μg/mL thiamine (Sigma-Aldrich), 0.4% glycerol (PanReac Applichem) with proper antibiotic at 37 °C with shaking at 200 rpm (Eppendorf Innova® 40). The overnight cultures were used to inoculate M9 glycerol medium to a starting OD₆₀₀ of 0.05. The culture was grown in an incubator at 37°C with shaking at 200 rpm until an OD₆₀₀ of 1.0. For the LS group, miliQ water was added to the suspension with a ratio of 46 μL per mL of culture, 1 mL of the suspension was kept aside for RNA isolation, 9 mL was used for 3C-based studies, 10 mL used for ChIP-based study. For the salt shock group, 5M NaCl was added to the suspension with a ratio of 46 μL per mL of culture resulting in a final concentration of 300 mM NaCl, 1 mL of the suspension was kept aside for RNA isolation, 9 mL was used for 3C-based studies, 10 mL used for ChIP-based study.

### RNA isolation

Total RNA was isolated from bacterial culture using PureLink® RNA Mini Kit (Catalog numbers: 12183020, Thermo Fisher scientific) according to the manufacturer’s protocol of purification from bacterial cells. The RNA concentration was measured by using NanoDrop™ 2000 spectrophotometer (Catalogue number: ND-2000, Thermo Scientific™) and NanoDrop 2000/2000c Software Version 1.6. Analyzing the quality of RNA sample by running 1% gel-red pre-stained agarose gel.

### RT-qPCR

#### Primer design

All primers **(Supplementary table1)** used for RT-qPCR were designed based on the *Escherichia coli* K-12 MG1655 reference genome (Accession number: NC_000913.3 – https://www.ncbi.nlm.nih.gov/nuccore/556503834) and were synthesized by Sigma-Aldrich (In solution, 100 μM). Primer specificity was confirmed by melting curve analysis.

#### Reaction set-up

RT-qPCR experiments were performed as described previously (18).

#### Data analysis

Data analysis was performed as described previously (18). To ensure comparability of gene expression levels, raw data are typically normalized using reference genes with relatively stable expression. According to RT-qPCR guidelines (98), at least three reference genes should be used to ensure data reliability. In this study, *dnaA*, *rpoD* and *rrsA* were used as reference genes. *dnaA* encodes the chromosomal replication initiator protein DnaA. *RpoD* encodes the RNA polymerase σ^70^ factor. *DnaA* and *rpoD* both have been used as reference control for RT-qPCR experiments in *E. coli* in different pH (99, 100) and osmolarity (18). *RrsA* codes for 16S rRNA. It has been used to normalize relative transcript levels in RT-qPCR studies of the *proVWX* operon (101). In the Results and Discussion sections of this study, all RT-qPCR data were normalized using *dnaA* as the reference control.

### ChIP-qPCR (102)

#### ChIP library preparation

Strains NT796, NT797, NT866 and NT867 (Table 1) were used for ChIP-qPCR. After bacterial growth under different pH conditions, the ChIP library was prepared as described in our earlier publication (102). To achieve high-resolution protein-binding profiles, the DNA fragment size in the ChIP library was carefully controlled within the range of 200–400 bp. Fragment size distribution was assessed by electrophoresis on a 1% gel-red pre-stained agarose gel.

#### Primer design

All primers **(Supplementary table2)** used for ChIP-qPCR were designed on the *Escherichia coli* K-12 MG1655 reference genome (Accession number: NC_000913.3 – https://www.ncbi.nlm.nih.gov/nuccore/556503834) and were synthesized by Sigma-Aldrich (In solution, 100 μM). Primer specificity was confirmed by melting curve analysis.

#### Reaction set-up

ChIP-qPCR experiments were performed as described previously (102).

#### Data analysis

Data analysis was performed as described previously (102). The binding efficiency of H-NS/StpA on *hdeD*1, *dnaA* and *appB* region were used for data normalization. Among them, *dnaA* has previously been used as a reference locus to map H-NS binding profiles in *Edwardsiella piscicida* by ChIP-qPCR (22). *appB* gene region is a H-NS and StpA negative binding region (23). *hdeD*1 produced data with smaller error bars after normalization. Therefore, data normalized using *hdeD*1 are presented in the figures. Equivalent data normalized with binding efficiency at *dnaA* have been provided in Supplementary figure 5e-h and Supplementary figure 9 e-h for comparison.

### 3C-qPCR (103, 104)

#### 3C library preparation

*E. coli* cells were harvested by centrifugation after cultivation under experimental conditions. Preparation of 3C libraries was performed as previously described (104). Notably, the restriction enzyme MluCI was used to digest the chromosome in all 3C studies of the *hdeAB* operon and its flanking regions, while NlaIII (18) was used for the *proVWX* operon and its flanking regions. Library quality was assessed by electrophoresis on a 1% gel-red pre-stained agarose gel.

#### Control library preparation

A synthetic control library was prepared for 3C-PCR **(supplementary table 3)**. The synthetic control template was generated by individually amplifying the chimeric fragments of interest by PCR and pooling them at equimolar ratios. This control library was then serially diluted to produce the standard samples for 3C-qPCR. The sequences of the synthetic control fragments for *proVWX* operon was same as previous study (18).

#### Primer and TaqMan probe design

All primers **(supplementary table 4)** used for 3C-qPCR were designed on the *Escherichia coli* K-12 MG1655 reference genome (Accession number: NC_000913.3 – https://www.ncbi.nlm.nih.gov/nuccore/556503834) and ordered from Sigma-Aldrich (In solution, 100 μM). The HPLC-purified, double-quenched TaqMan probes **(supplementary table 5)** were designed as described previously (103), which is with a 5’ 6-FAM fluorophore, 3’ Iowa Black™ Fluorescence Quencher and an internal ZEN quencher were ordered from Integrated DNA Technologies. All of the primers and probes were dissolved in 1X TE to a final concentration of 100 μM and stored at −20°C.

#### Reaction set-up and data analysis

The PrimeTime™ Gene Expression Master Mix Kit (Catalogue number: 10557710, Integrated DNA Technologies, Inc.) was used for 3C-qPCR. The experiments reaction set-up and data analysis was performed as described previously (18, 104).

### Data visualization and statistical analyses

Data visualization for RT-qPCR, ChIP-qPCR and 3C-qPCR studies and the associated statistical analyses were performed by using GraphPad Prism (version 10.1.0). Statistical significance was determined by unpaired, two-tailed Student’s t-test. P-values indicate the level of significance: *p* < 0.05 (*), *p* < 0.01 (**), *p* < 0.001 (***) and ns indicates no significant difference.

### Bacterial spotting assay

A single bacterial colony was used to inoculate 2 mL of M9 medium. The cells were cultured overnight at 37 °C with shaking at 200 rpm. The overnight culture was then used to inoculate 5 mL of fresh M9 medium to an initial OD₆₀₀ of 0.01. Cultures were grown up to an OD₆₀₀ of 1.0. Cells were then diluted with fresh M9 medium to an OD₆₀₀ of 0.5. Serial 10-fold dilutions of the cell suspension were prepared in Eppendorf tubes. For the acid stress group, the pH of the diluted cell suspension was adjusted to 2.0 using 37% HCl and cells were incubated at room temperature for 30 minutes. After incubation, 10 μL of the treated (pH 2.0) and control (pH 6.6) cell suspensions were spotted onto M9 agar plates and incubated overnight at 37 °C. The plates were imaged using the Bio-Rad universal hood II gel doc imaging system.

### Genomic DNA isolation

A single *E. coli* (NT331) colony was cultured in 2 mL LB medium up to stationary phase at 37 °C with shaking at 200 rpm. Genomic DNA was isolated as previously described (18). Genomic DNA was dissolved in nuclease-free water. The DNA concentration was measured using the Qubit® dsDNA HS Assay Kit (Catalogue number: Q32854, ThermoFisher Scientific).

### H-NS cloning and purification

H-NS protein was purified as described previously (18). H-NS was dissolved in storage buffer (20 mM Tris pH 7.5, 300 mM KCl, 10% glycerol and β-mercaptoethanol) and stored at −80 °C.

### Microscale thermophoresis (MST)

H-NS protein was serially diluted from 54.6 to 0.0016 μM using either pH 7.5 dilution buffer (20 mM Tris–HCl, pH 7.5, 300 mM KCl, 10% glycerol, 0.1% Tween-20) or pH 5.0 dilution buffer (20 mM sodium acetate, pH 5.0, 300 mM KCl, 10% glycerol, 0.1% Tween-20). Each dilution (10 μL) was then mixed 1:1 with 80 nM DNA substrate (Cy5 labeled, 78 bp, 32%GC content) prepared in Milli-Q water, yielding a final DNA concentration of 40 nM and H-NS concentrations ranging from 27.3 to 0.0008 μM in measurement buffer (10 mM Tris–HCl, pH 7.5, or 10 mM sodium acetate, pH 5.0, 150 mM KCl, 5% glycerol, 0.05% Tween-20). Samples were loaded into MST capillaries (Monolith NT.115 Standard Capillaries, NanoTemper Technologies, Germany), and measurements were performed using a Monolith NT.115 instrument (NanoTemper Technologies) at 40% LED power and medium MST power. Each measurement consisted of a total acquisition time of 40 s, including 5 s laser off, 30 s laser on, and 5 s laser off. Data values were evaluated after 20 seconds of laser on. Data analysis by using equipment attached software (MO. Affinity Analysis v2.1.5) and plot by using GraphPad Prism (version 10.1.0).

### DNA bridging assay

DNA bridging assays were performed as previously described, with minor modifications (14). Briefly, 57 μL of streptavidin-coated paramagnetic Dynabeads M-280 (Invitrogen) were washed once with 100 μL 1× PBS and twice with coupling buffer (20 mM Tris–HCl, pH 8.0, 2 mM EDTA, 2 M NaCl, 2 mg/mL acetylated BSA, and 0.04% Tween-20), according to the manufacturer’s instructions. After washing, beads were resuspended in 57 μL coupling buffer and divided into two aliquots (39 μL and 18 μL). To the 39 μL bead suspension, 39 μL buffer (10 mM Tris–HCl, pH 8.0, 20 mM KCl, 20 mM MgCl₂) containing biotinylated bait DNA (44 ng per sample) was added. The 18 μL bead suspension received 18 μL buffer (10 mM Tris–HCl, pH 8.0, 20 mM KCl, 20 mM MgCl₂) alone. Samples were incubated at 25 °C for 20 min with shaking at 1000 rpm to allow bait DNA immobilization. Following incubation, beads were washed twice with incubation buffer (10 mM Tris–HCl, pH 8.0, 50 mM KCl, 10 mM MgCl₂, 5% (v/v) glycerol, 1 mM DTT, and 1 mM spermidine) using volumes of 182 μL or 84 μL, corresponding to the initial bead aliquots. Beads were then resuspended in the same volumes of incubation buffer and mixed with 26 μL or 12 μL of prey DNA (^32^P-labeled, 685 bp, 32% GC content) together with H-NS protein at the indicated concentrations. Reactions were incubated at 25 °C for 20 min with shaking at 1000 rpm. To remove unbound prey DNA, beads were washed using wash buffer and subsequently resuspended in 12 μL stop buffer (10 mM Tris–HCl, pH 8.0, 1 mM EDTA, 200 mM NaCl, and 0.2% SDS). Samples were transferred to PCR tubes, and recovered radioactivity was quantified by liquid scintillation counting. All values obtained from DNA bridging assays were corrected for background signal using control samples lacking H-NS and normalized to a reference sample containing the total amount of ^32^P-labeled prey DNA used in the assay. DNA bridging experiments were performed in duplicates.

## Results

### Deletion of *hns* and *stpA* leads to upregulation of acid resistance genes, enhancing survival under acid stress

H-NS controls the cellular acid resistance system by repressing the transcription of genes involved in the acid response under uninduced conditions (29). To examine the importance of H-NS and StpA in repression of these genes, we constructed Δ*hns,* Δ*stpA* and Δ*hns* Δ*stpA* strains **(Table 1)** and examined their resistance to acid in a spotting assay on M9 agar plates at pH 6.6 and pH 2.0 (see Materials and Methods). As expected, the H-NS-deficient strains exhibited significantly enhanced acid resistance at pH 2.0 **(Figure 1a, b)**. In the *stpA* single mutant strain, only a mild increase in acid resistance is observed compared to the WT strain **(Figure 1a, b)**, which is because of repression of genes for acid resistance mediated by H-NS. Overall, the enhanced acid resistance observed in *E. coli* strains lacking H-NS can be attributed to alleviation of H-NS-mediated transcription repression of acid response genes.

**Figure 1.**
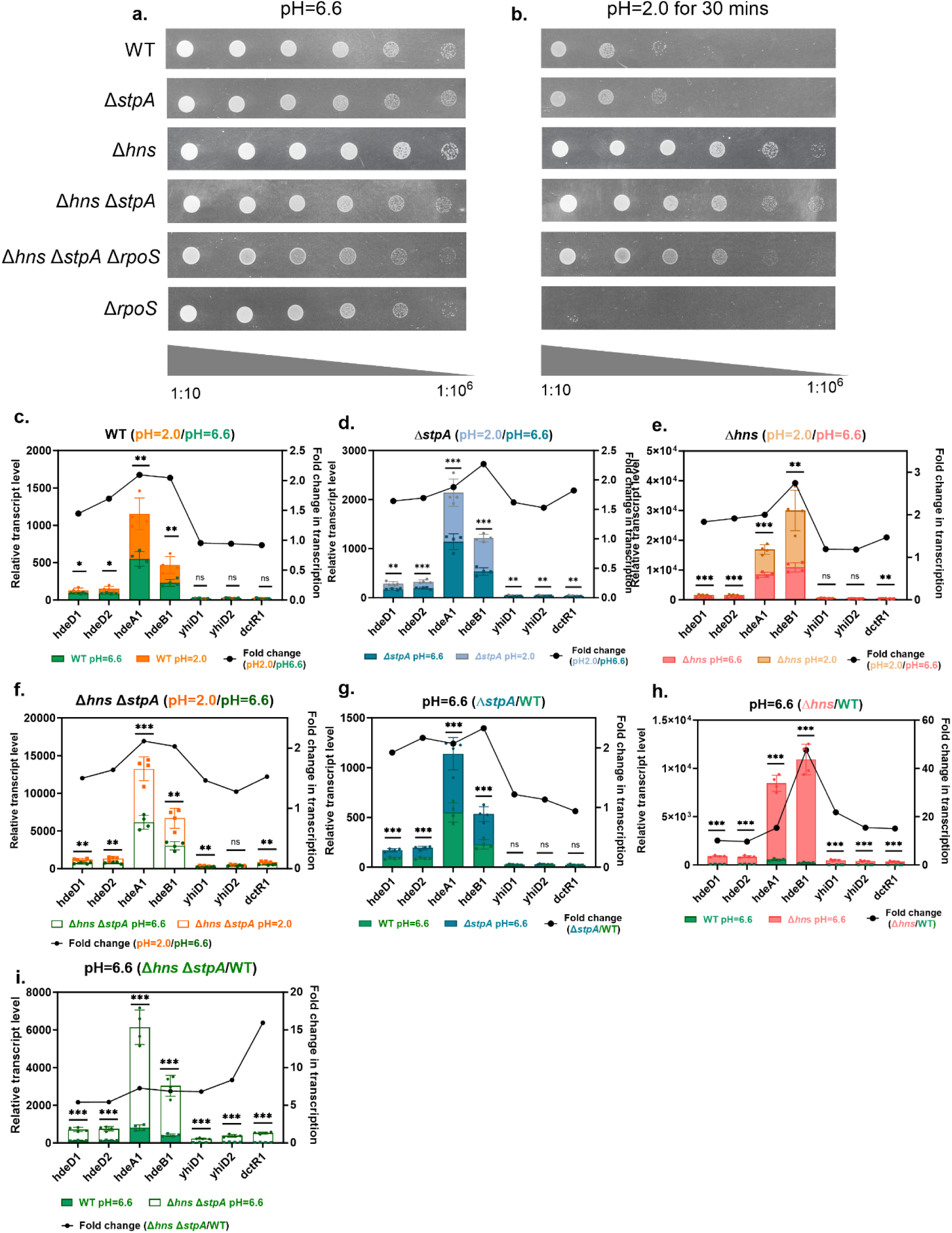
Deletion of *hns* and *stpA* leads to upregulation of acid resistance genes, enhancing survival under acid stress. **a–b.** Spotting test of WT, Δ*stpA,* Δ*hns*, Δ*hns* Δ*stpA*, Δ*hns* Δ*stpA* Δ*rpoS* and Δ*rpoS* strains on M9 plates at pH 6.6 (**a**) and pH 2.0 (**b**). Serial 10-fold dilutions are spotted from left to right. Transcription profile of the *hdeAB* operon and its flanking genes at pH 6.6 and pH 2.0 conditions in the WT (**c**), Δ*stpA* (**d**), Δ*hns* (**e**) and Δ*hns* Δ*stpA* (**f**) strains. Transcription profile of *hdeAB* operon and its flanking genes at pH 6.6 in the Δ*stpA* (**g**), Δ*hns* (**h**) or Δ*hns* Δ*stpA* (**i**) strains compared with WT. Bar graphs (follow left y-axis) represent relative transcript levels; line graphs (follow right y-axis) indicate fold changes in gene transcription between conditions or strains. Each group includes four biological replicates and three technical replicates. *dnaA*, *rpoD* and *rrsA* were tested as internal reference genes for normalization, yielding similar results; *dnaA* is used for data presentation. Statistical significance was determined by unpaired, two-tailed Student’s *t*-test. *p*-values indicate the level of significance: *p* < 0.05 (\**), p < 0.01 (**), p < 0.001 (\*\*\**) and *ns* indicates no significant difference.

To confirm and extend observations that H-NS and StpA mediate repression of acid resistance genes, we used RT-qPCR to generate a transcription profile for the acid response operon *hdeAB* in *E. coli* at exponential phase. We designed seven pairs of primers for this operon and its flanking genes **(Supplementary table 1)**, the genes flanking the *hdeAB* operon also belong to the AFI **(Supplementary figure 2)** but the exact functions of their gene products are still unknown (54, 105). All strains were cultured in supplemented M9 glycerol medium and grown to an OD_600_ of ∼1.0 (exponential phase). To validate the applied acid shock conditions, preliminary experiments were conducted to assess the transcription profile of *hdeAB* following incubation at pH 4.0 for 30 minutes and pH 2.0 for 5 and 10 minutes. The transcription of *hdeAB* was promptly activated after only 5–10 minutes of exposure to the extreme acidic condition (pH 2.0). In contrast, no upregulation of *hdeAB* transcription was observed after 30 minutes under mildly acidic conditions (pH 4.0) **(Supplementary figure 3a)**. Based on these findings, all of the experiments with acid shock group cells were conducted using a 10 minutes incubation at pH 2.0 **(Supplementary figure 3b)**. RT-qPCR shows that the transcription of *hdeA* and *hdeB* is activated by acid shock and the transcription level increases 2-fold upon acid shock in wild type (WT) **(Figure 1c)**. The transcription of the upstream gene *hdeD* also increases under acid stress, while the transcription level of downstream *yhiD* and *dctR* genes remains unaltered **(Figure 1c)**. We also tested the transcription of these genes in the Δ*hns*, Δ*stpA* and double mutant (Δ*hns* Δ*stpA*) strains. The transcription of *hdeD*, *hdeA* and *hdeB* was activated upon acid shock in all these strains **(Figure 1d-f)**. These observations confirm that the transcription of *hdeAB* operon is induced by acid stress, which is in agreement with previous studies (75, 105). Interestingly, upon acid shock, the transcription of *yhiD* and *dctR,* positioned downstream of the *hdeAB* operon, also exhibited a significant ∼1.5-fold increase in the Δ*hns,* Δ*stpA* and Δ*hns* Δ*stpA* strains **(Figure 1d-f)**. Compared to the WT strain, deletion of *stpA* alone resulted in a modest upregulation (2-fold) of *hdeD*, *hdeA* and *hdeB* only, while the transcript levels of the downstream genes *yhiD* and *dctR* remained unaffected **(Figure 1g)**. Deletion of *hns* yielded a much larger increase in the transcription levels of the detected amplicons (at least 10-fold), with *hdeA* and *hdeB* showing particularly strong upregulation-exceeding 20-fold and 30-fold **(Figure 1h)**. As expected, the double mutant strain of StpA and H-NS also resulted in a substantial increase in transcription levels, ranging from 5- to 20-fold **(Figure 1i)**. Taken together, these data indicate an important regulatory role for H-NS and StpA, with minor effects observed in the *stpA* single deletion strains, possibly attributable to the cross-regulation between H-NS and StpA and a lower expression level of StpA in the presence of H-NS **(Supplementary figure 4a).**

### H-NS binding at the *hdeAB* promoter is reduced under acidic pH conditions

Is the observed repression described in the previous section attributed to the direct binding of H-NS and StpA? To answer this question, we used chromatin immunoprecipitation-quantitative PCR (ChIP-qPCR) to examine the binding profiles of H-NS and StpA to the *hdeAB* operon and its flanking regions at different pH conditions **(Figure 2)**. Based on a previous genome wide ChIP-chip study of H-NS and StpA (23), we constructed strains expressing H-NS and StpA with a 3X FLAG-tag at the C-terminus **(Table 1)**. To obtain reliable ChIP-qPCR results, the binding efficiency of H-NS and StpA, respectively at three different regions (*hdeD*1, *dnaA* or *appB*) was used for ChIP-qPCR data normalization **(see Materials and Methods)**. ChIP-qPCR analysis revealed that H-NS has a broader range of enrichment across our target region **(Figure 2a, e)**, consistent with a previous ChIP-chip analysis (23). Binding of H-NS in this region suggests that H-NS directly regulates the *hdeAB* operon (as observed in our RT-qPCR experiments). At acid shock (pH=2.0), H-NS exhibited a reduced binding signal at the *hdeAB* promoter region **(Figure 2a, b)**. With different internal controls used for data analysis, the fold change of H-NS relative binding efficiency (pH2.0/pH6.6) exhibited a similar trend **(Figure 2b)**, establishing the reliability of the ChIP-qPCR results. To eliminate the potential influence of StpA, we also examined the binding profile of H-NS under different pH conditions in a Δ*stpA* strain. In this strain, H-NS binding exhibited similar sensitivity to acidic conditions **(Supplementary figure 5a, b)**. Next, we examined the binding of StpA and found that its association with the region surrounding the operon was minimally affected by pH changes **(Figure 2c, d)**. However, in the absence of H-NS, a reduced binding efficiency of StpA under acidic conditions became more pronounced **(Supplementary figure 5c, d)**. Since H-NS and StpA form functional heterodimers, we next investigated whether H-NS or StpA affects the binding ability of its counterpart. To this end, we separately compared the H-NS binding in strains with and without StpA and the StpA binding signal in strains with and without H-NS. The results indicated that StpA does not affect H-NS binding at the *hdeAB* operon **(Figure 2e, f)**. However, deletion of H-NS leads to a significant increase in StpA binding at the *dctR* locus, while no detectable change in StpA binding was observed at the *hdeAB* promoter region **(Figure 2g, h)**.

**Figure 2.**
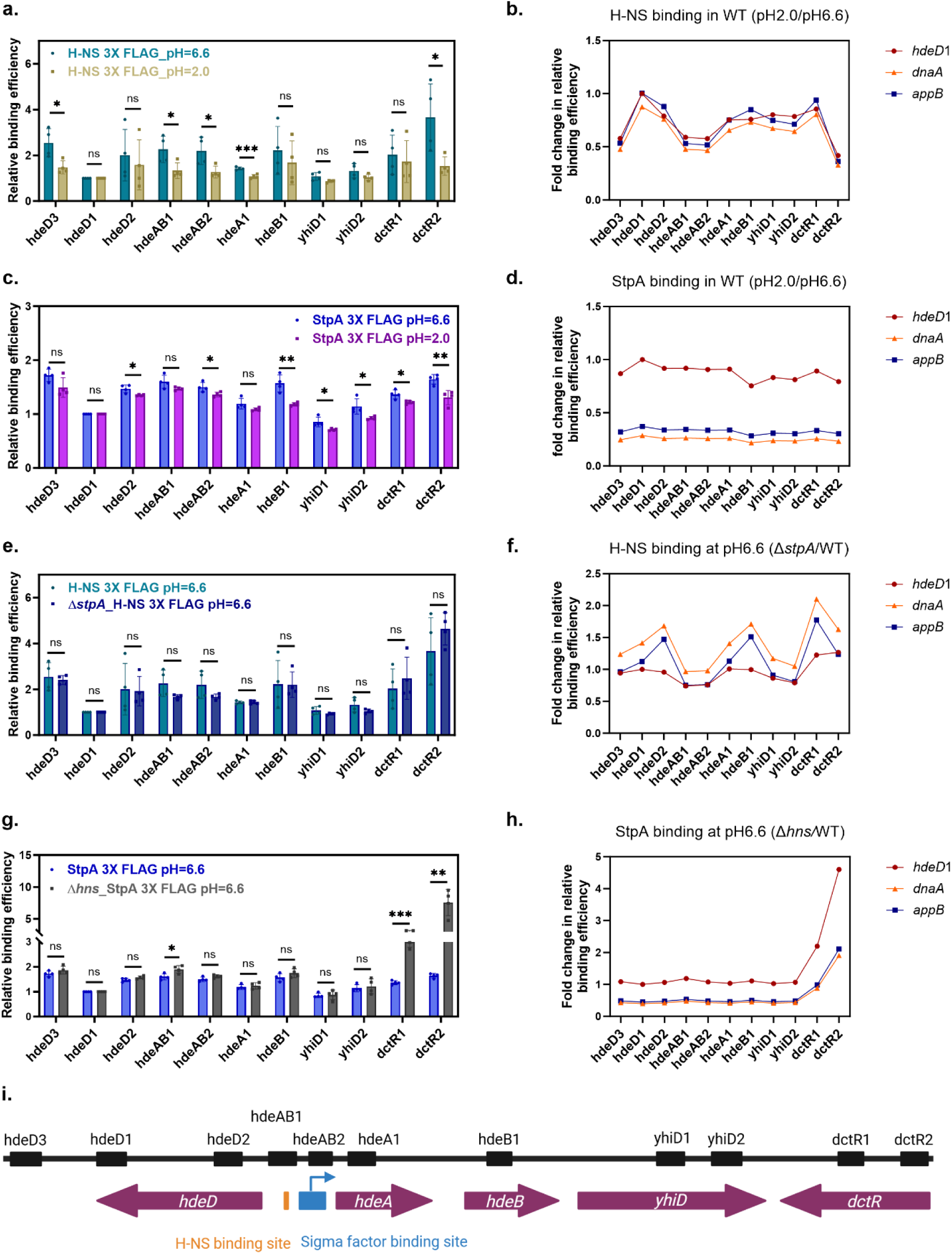
Binding profiles of H-NS and StpA at the *hdeAB* operon and its flanking genes. **a.** Binding profile of H-NS at *hdeAB* operon and its flanking genes at pH 6.6 and pH 2.0 conditions. **b.** Fold change of H-NS relative binding efficiency (pH2.0/pH6.6) with different reference loci, *hdeD, dnaA,* and *appB*, respectively. **c.** Binding profile of StpA at *hdeAB* operon and its flanking gene at pH 6.6 and pH 2.0 conditions. **d.** Fold change of StpA relative binding efficiency (pH2.0/pH6.6) with controls as in b. **e.** Binding profile of H-NS at *hdeAB* operon and its flanking genes at pH 6.6 in the presence and absence of StpA. **f.** Fold change of H-NS relative binding efficiency (Δ*stpA*/WT) with controls as in b. **g.** Binding profile of StpA at *hdeAB* operon and its flanking gene at pH 6.6 in the presence and absence of H-NS. **h.** Fold change of H-NS relative binding efficiency (Δ*hns*/WT) with different controls as in b. **i.** Schematic showing the positions of amplicons used in ChIP-qPCR analysis. Each group includes four biological replicates with two technical replicates each. The binding efficiency of H-NS/StpA at *hdeD*1, *dnaA* and *appB* were used for data normalization. The presented data are using *hdeD*1 as the reference. Statistical significance was determined by unpaired, two-tailed Student’s *t*-test. *p*-values indicate the level of significance: *p* < 0.05 (\**), p < 0.01 (**), p < 0.001 (\*\*\**) and *ns* indicates no significant difference.

### *hdeAB* transcription is regulated by H-NS-mediated chromatin remodeling

H-NS and StpA have been shown to organize DNA by forming DNA bridges and filament-like structures, thereby contributing to the formation of functionally organized, compact chromatin (12, 56). Given the changes in H-NS and StpA binding at the *hdeAB* promoter region upon acid shock, we hypothesized that these changes may lead to local chromatin remodeling. To test this hypothesis, we employed chromosome conformation capture-qPCR (3C-qPCR) and ‘visualized’ the local chromatin architecture in this region. 3C-qPCR is a technique used to quantify the physical interactions between an anchor fragment and target genomic regions, thereby revealing three-dimensional DNA organization within the genome. The method relies on formaldehyde crosslinking to preserve chromatin interactions, followed by restriction enzyme digestion and intramolecular ligation. Quantitative PCR is then used to measure the frequency of ligation products, which correlates with the spatial proximity of genomic loci *in vivo* (see Materials and Methods). In this case, we used the restriction enzyme MluCI to digest the target genomic region. This enzyme efficiently fragments the region into relatively uniform fragments of appropriate size for subsequent qPCR detection **(Figure 3)**. We used fragment hdeAB10_MluCI (red colored number at x-axis in figure 3) that contains the sigma factor binding site of the *hdeAB* operon as an anchor fragment and determined its relative interaction frequency with other restriction digestion fragments of and around the *hdeAB* operon.

**Figure 3.**
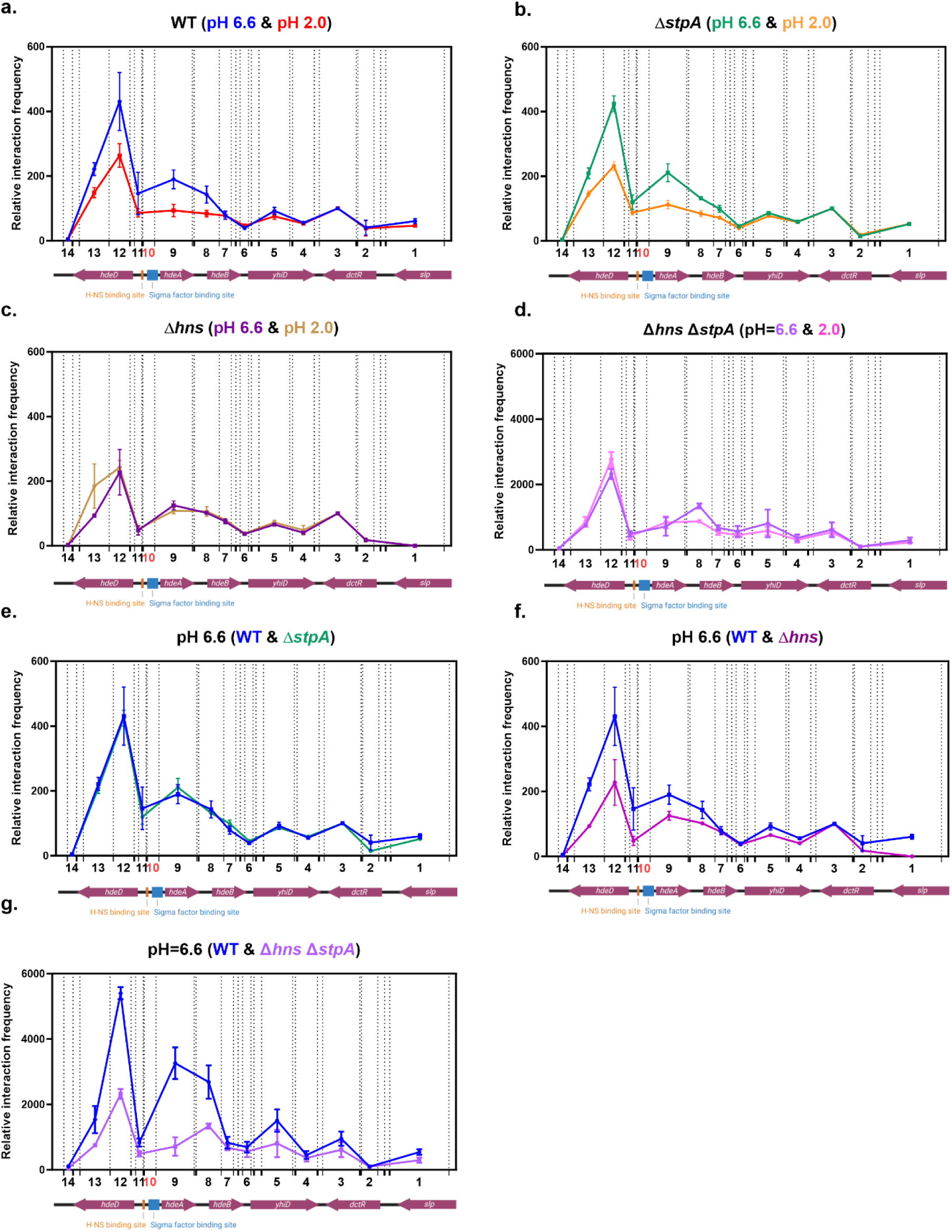
Local chromatin architecture of the *hdeAB* operon and its flanking genes. Relative interaction frequency at the *hdeAB* operon and its flanking genes in WT (**a**), Δ*stpA* (**b**), Δ*hns* (**c**) and Δ*hns* Δ*stpA* (**d**) strains at pH 6.6 and pH 2.0. Relative interaction frequency of the *hdeAB* operon and its flanking genes at pH 6.6 condition in Δ*stpA* (**e**), Δ*hns* (**f**) or Δ*hns* Δ*stpA* (**g**) strains compared with WT. Dashed lines indicate the MluCI restriction enzyme cleavage sites within the *hdeAB* operon. The x-axis shows the relative positions and lengths of target fragments and the anchor fragment (fragment 10, highlighted in red) used in this study. The y-axis represents the relative interaction frequency between the anchor and target fragments.

3C-qPCR analysis shows the fragment 12 has the highest relative interaction frequency with anchor fragment (fragment 10), which indicates a hairpin-like chromatin structure maintained at the regulatory region of *hdeAB* operon **(Figure 3a)**. As described above, a significant upregulation of *hdeAB* transcription in the WT and Δ*stpA* strain was observed after acid shock (pH 2.0) **(Figure 1c-d)**, accompanied by a reduction in H-NS binding at the operon’s promoter region **(Figure 2a, supplementary figure 5a)**. Consistently, a clear decompaction of the chromatin structure upstream of and across the *hdeAB* operon (fragment 8 to 13) was observed following acid treatment **(Figure 3a-b)**. Collectively, these findings demonstrate a correlation between transcription activation of the operon and local chromatin remodeling. To further determine the requirement of H-NS and StpA for maintaining this hairpin-like chromatin structure, we compared chromatin architecture between mutant and WT strains. In the Δ*stpA* strain, H-NS remains bound as a homodimer to maintain chromatin organization **(Figure 3e)**. Correspondingly, only a modest increase in transcription of the *hdeAB* operon and its flanking genes was observed **(Figure 1d)**. In contrast, deletion of *hns* disrupted chromatin architecture **(Figure 3f-g)**, underlining the structural role of H-NS in this region. StpA was unable to act as backup. This disruption relieved H-NS-mediated transcription repression, as evidenced by the dramatic increase in *hdeAB* transcription observed in the Δ*hns* and Δ*hns* Δ*stpA* strains **(Figure 1h-i)**. In the *hns* mutant and *hns* Δ*stpA* double mutant strains, the chromatin conformation near the *hdeAB* region was not affected by acid shock **(Figure 3c-d)**. This is attributed to the absence of the primary structural protein, H-NS, resulting in an open chromatin conformation. However, the increased transcription of *hdeAB* after acid shock in *hns* mutants and *hns stpA* double mutant **(Figure 1e-f)**, suggests the involvement of additional regulatory factors in the transcriptional response to acid shock of the *hdeAB* operon.

We have established that H-NS responds to acid shock by remodeling local chromatin conformation and relieve of transcriptional repression of the *hdeAB* operon. As shown in previous studies, the sigma factors RpoD and RpoS competitively bind at the same site and recruit the core RNA polymerase to form the holoenzyme for transcription of *hdeAB* (71, 76). Shin et al. demonstrated that only RpoD-mediated transcription is repressed by H-NS (71). Our RT-qPCR analysis revealed a significant increase in *rpoS* transcript levels following acid shock **(Supplementary figure 4b)**. But it is not clear in what differences at the protein level it translates at this time scale. To further investigate the H-NS-dependent transcriptional regulation of this operon, we constructed Δ*rpoS* mutants and Δ*hns* Δ*stpA* Δ*rpoS* triple mutants **(Table 1)** to exclude secondary effects of RpoS. This ensures that transcription under our experimental conditions is driven by RpoD only and that any measured changes in our system are not controlled via the RpoS-dependent general stress response. We then assessed *hdeAB* transcript levels and characterized the local chromatin conformation in these strains.

The RT-qPCR data show that the transcription of *hdeAB* and its flanking genes increased upon acid shock in the Δ*rpoS* strain **(Figure 4a)**. Compared to WT, the transcription of all amplicons sharply decreased in the absence of RpoS. Transcription of *hdeA* and *hdeB* decreased by about 75% **(Figure 4b)**. This indicates that under non-acidic conditions during the exponential phase, the levels of RpoS protein are sufficient to drive part of the transcription of *hdeAB.* This finding is supported by high-throughput transcriptomic analyses revealing that RpoS participates in the transcriptional regulation of an expanding set of genes during the exponential phase of growth (106). These observations contradict the classical view that RpoS plays only a minor role in recruiting RNA polymerase in this growth phase, attributed to its relatively low expression level compared with that in the stationary phase (107, 108). Consistent with the RT-qPCR results, an open chromatin conformation was observed at the *hdeAB* operon locus in the Δ*rpoS* mutant upon acid shock **(Figure 4e)**. This reaffirms our model that the dynamic structural changes in this region are driven by a pH-specific response of H-NS. What stands out is that the absence of RpoS decreases the anchor interaction frequency with fragment 8 **(Figure 4f)**.

**Figure 4.**
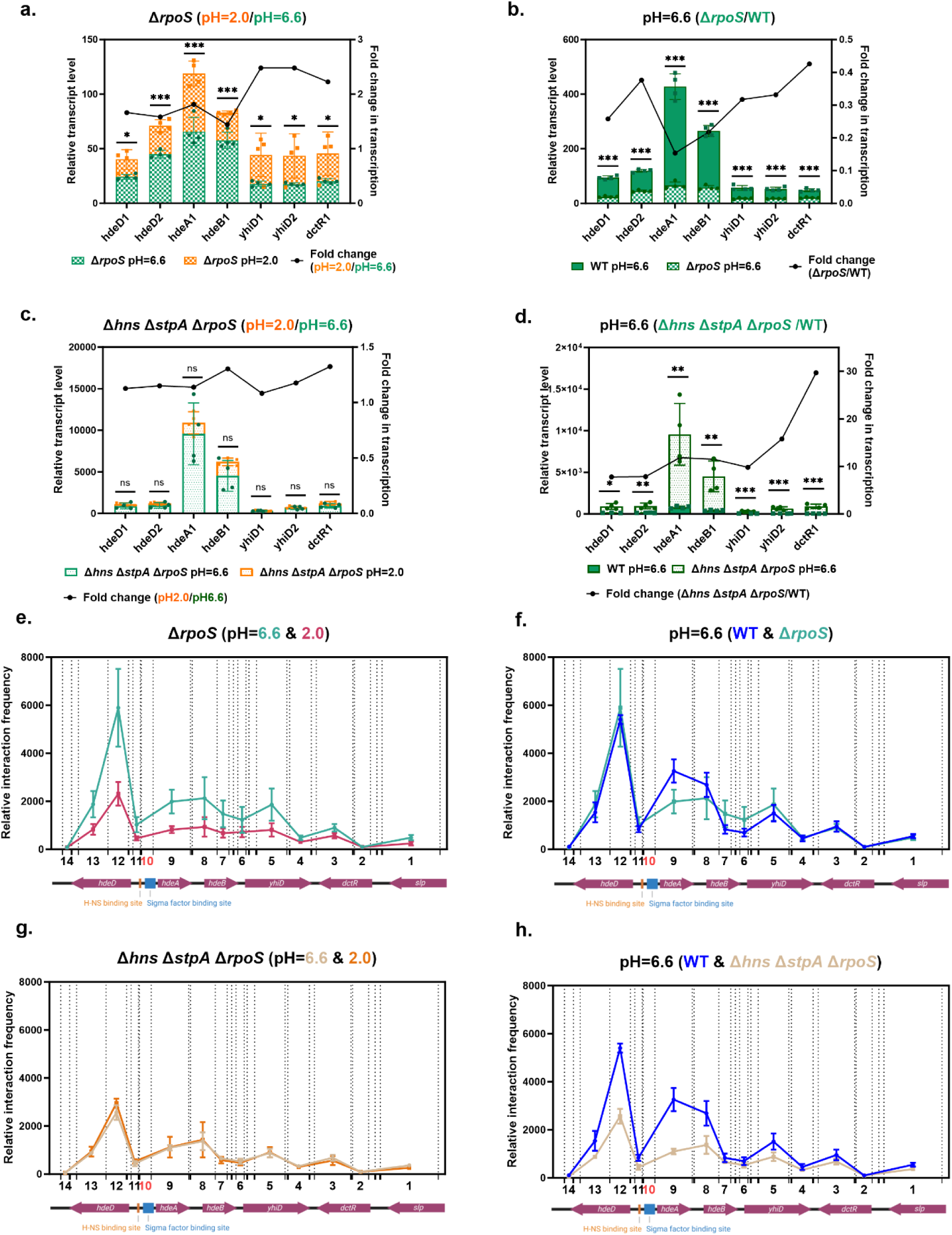
Transcription profiles and local chromatin architecture of the *hdeAB* operon and its flanking genes in Δ*rpoS* and Δ*hns* Δ*stpA* Δ*rpoS* strains. Transcription profile of the *hdeAB* operon and its flanking genes at pH 6.6 and pH 2.0 conditions in the Δ*rpoS* (**a**) and Δ*hns* Δ*stpA* Δ*rpoS* (**c**) strains. Transcription profile of *hdeAB* operon and its flanking genes at pH 6.6 in the Δ*rpoS* (**b**) or Δ*hns* Δ*stpA* Δ*rpoS* (**d**) strains compared with WT strain. Bar graphs (follow left y-axis) represent relative transcript levels; line graphs (follow right y-axis) indicate fold changes in gene transcription between conditions or strains. Each group includes four biological replicates and three technical replicates. *dnaA*, *rpoD* and *rrsA* were tested as internal reference genes for normalization, yielding similar results; *dnaA* is used for data presentation. Statistical significance was determined by unpaired, two-tailed Student’s *t*-test. *p*-values indicate the level of significance: *p* < 0.05 (\**), p < 0.01 (**), p < 0.001 (\*\*\**) and *ns* indicates no significant difference. Relative interaction frequency at *hdeAB* operon and its flanking genes in the Δ*rpoS* (**e**) and Δ*hns* Δ*stpA* Δ*rpoS* (**g**) strain at pH 6.6 and pH 2.0 conditions. Relative interaction frequency of *hdeAB* operon and its flanking genes at pH6.6 condition in Δ*rpoS* (**f**), Δ*hns* Δ*stpA* Δ*rpoS* (**h**) compared with WT. The x-axis shows the relative positions and lengths of target fragments and the anchor fragment (fragment 10, highlighted in red) used in this study. The y-axis represents the relative interaction frequency between the anchor and target fragments.

Compared to the acid-induced transcription activation of *hdeAB* observed in the RT-qPCR experiments on single and double mutant strains, in the Δ*hns* Δ*stpA* Δ*rpoS* strain, the acid-dependent transcription activation of the *hdeAB* operon was abolished **(Figure 4c)** and the local chromatin structure was also not affected by acid shock, which indicated the loss of pH-dependent structural remodeling in the absence of these regulators **(Figure 4g)**. Compared to the WT strain, the transcription levels of *hdeAB* and its flanking genes remain substantially elevated (5- to 30-fold) in the Δ*hns* Δ*stpA* Δ*rpoS* strain **(Figure 4d)**. Due to the absence of H-NS and StpA, chromatin structure is ‘open’ and the promoter region is consistently exposed **(Figure 4h)**. A recent Micro-C chromatin conformation capture study revealed that H-NS and StpA can cooperatively form chromosomal hairpin-like structures that repress transcription of nearby genes (109). To validate our proposed model in which H-NS forms a hairpin-like structure at the *hdeAB* promoter region, we visualized Micro-C data spanning the *hdeAB* operon and its flanking 5 kb regions upstream and downstream. We observed prominent vertical clusters of contacts at the *hdeAB* promoter, a pattern that directly indicates the presence of a hairpin-like structure (**Supplementary figure 6**). Notably, this structure was preserved in the *stpA* deletion strain. In contrast, the hairpin structure was completely disrupted in the *hns* deletion strain and in the *hns stpA* double mutant, where it was replaced by an operon-sized chromosomal interaction domain (OPCID). Such domains have been associated with high transcriptional activity (109), and is supported by our RT-qPCR data **(Figure 1h, i)**.

To further investigate whether the reduced DNA-binding capacity of H-NS following acid shock *in vivo* results from a direct response of H-NS to acidic conditions (**Figure 2a**), we examined the DNA-binding affinity of purified H-NS protein under different pH conditions using microscale thermophoresis (MST). Previous studies have shown that when *E. coli* is exposed to an external pH of 2.0, its intracellular pH remains close to 4.0 (110). Accordingly, we initially attempted to measure H-NS DNA-binding at pH 4.0. However, protein aggregation was observed under these conditions (data not shown). We therefore increased the experimental pH to 5.0. Compared with neutral pH conditions (pH=7.5), H-NS exhibited a significantly reduced DNA-binding ability under acidic conditions (pH=5.0) **(Supplementary Figure 7a)**. To investigate the effect of pH on the DNA-bridging activity of H-NS *in vitro*, we next employed the same experimental approach used previously to characterize H-NS-DNA bridging (14). In this assay, H-NS bridges biotin-labeled bait DNA immobilized on streptavidin-coated magnetic beads with ^32^P-labeled prey DNA. The resulting DNA–protein complexes were recovered by magnetic pull-down, and the radioactive signal of the recovered DNA served as a quantitative measure of DNA-bridging efficiency. The DNA-bridging efficiency of H-NS at pH=5.0 is markedly lower than that at pH=7.5 **(Supplementary Figure 7b)**. Together, these *in vitro* experiments provide direct evidence that H-NS responds directly to acidic environments, leading to impaired DNA binding and consequently to reduced DNA-bridging activity.

Collectively, these observations lend support to a model in which H-NS forms a repressive hairpin-like structure at the promoter region of the *hdeAB* operon, which under acidic conditions is ruptured by the reduced DNA-binding capacity of H-NS, resulting in acid-induced transcriptional activation of the *hdeAB* operon.

### The transcription of *proVWX* is not sensitive to pH fluctuation

In a previous study, we examined the role of H-NS in the osmotic regulation of the osmo-responsive *proVWX* operon (18). We showed that following a 10-minute salt shock from 0.08 M to 0.3 M NaCl, the transcription level of *proVWX* in *E. coli* exhibited a marked upregulation of 6- to 8-fold. Relief of repression by H-NS was due to remodeling of local chromatin architecture attributed to an ion-induced change in H-NS conformation and DNA binding mode (14, 18). Given that acid shock remodels the higher-order chromatin architecture maintained by H-NS at the *hdeAB* operon, thereby relieving its transcriptional repression, we next examined the effect of acid shock on *proVWX* transcription. This information is important in the light of response specificity. We examined the transcription of *proVWX* and its flanking genes at 0.17 M NaCl under different pH conditions in all bacterial strains described above. The transcription of *proVWX* is minimally responsive to acid shock: transcription of *proV* and *proW* in WT decreased by ∼25%, the transcription of *proX* is not affected by acid shock **(Figure 5a)**. The effect is more pronounced and extended across the operon in *stpA* and *hns* deletion mutants **(Supplementary figure 8a, b)**. Transcription across and flanking the *proVWX* operon was not affected by acid shock in Δ*rpoS*, Δ*hns* Δ*stpA* and Δ*hns* Δ*stpA* Δ*rpoS* strains **(Supplementary figure 8c-e)**. Comparison of WT and *stpA* deletion strains revealed that absence of StpA partially relieved transcription repression of the operon, specifically affecting *proV* and *proW*, while *proX* expression remained unchanged **(Figure 5b)**. On the other hand, H-NS exhibited a broader gene repressive effect across the entire *proVWX* operon. However, the magnitude of transcription upregulation in the *hns* deletion strain was modest, only 1.5- to 2.0-fold **(Figure 5c)**. A similar phenomenon was also observed in the Δ*hns* Δ*stpA* and Δ*hns* Δ*stpA* Δ*rpoS* strains, where upregulation of *proVWX* operon transcription was no more than 4-fold **(Figure 5d, e, supplementary figure 8)**, substantially lower than the level of repression H-NS exerts on *hdeAB*. By comparing the *rpoS* mutant strain with the WT, we found that the overall transcription level of *proVWX* increased by approximately 10% in the absence of RpoS, with the difference reaching statistical significance based on a t-test **(Figure 5f)**. Previous studies using β-galactosidase assays have shown that although *proVWX* harbors both RpoS and RpoD dependent promoters, *proVWX* transcription is not affected by the deletion of *rpoS* (111). By using other reference genes, we confirmed the lack of an outspoken effect **(Supplementary figure 8f)**.

**Figure 5.**
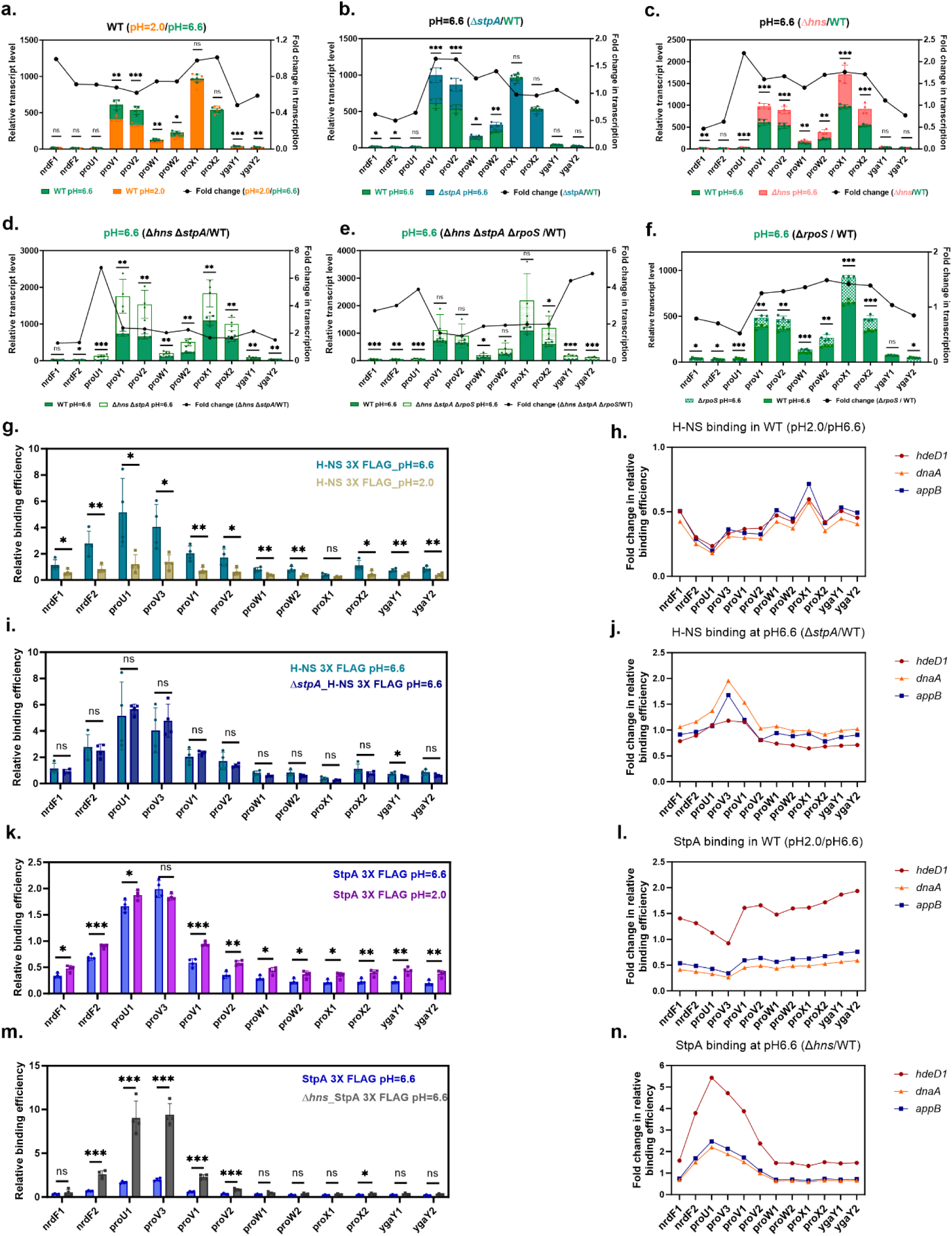
Regulation of the *proVWX* operon by H-NS and StpA. **a.** Transcription profiles of *proVWX* operon and its flanking genes at pH 6.6 and pH 2.0 conditions in the WT strain. Transcription profile of *proVWX* operon and its flanking genes at pH 6.6 in the Δ*stpA* (**b**), Δ*hns* (**c**), Δ*hns* Δ*stpA* (**d**), Δ*hns* Δ*stpA* Δ*rpoS* (**e**) and Δ*rpoS* (**f**) strains compared with WT. Bar graphs (follow left y-axis) represent relative transcript levels; line graphs (follow right y-axis) indicate fold changes in gene transcription between conditions or strains. Each group includes four biological replicates and three technical replicates. *dnaA*, *rpoD* and *rrsA* were tested as internal reference genes for normalization, yielding similar results; *dnaA* is used for data presentation. Statistical significance was determined by unpaired, two-tailed Student’s *t*-test. *p*-values indicate the level of significance: *p* < 0.05 (\**), p < 0.01 (**), p < 0.001 (\*\*\**) and *ns* indicates no significant difference. **g.** Binding profile of H-NS at *proVWX* operon and its flanking genes at pH 6.6 and pH 2.0 conditions. **h.** Fold change of H-NS relative binding efficiency (pH2.0/pH6.6) with different internal controls. **i.** Binding profile of H-NS at *proVWX* operon and its flanking genes at pH 6.6 in the presence and absence of StpA strains. **j.** Fold change of H-NS relative binding efficiency (Δ*stpA*/WT) with different internal controls. **k.** Binding profile of StpA at *proVWX* operon and its flanking genes at pH 6.6 and pH 2.0 conditions. **l.** Fold change of StpA relative binding efficiency (pH2.0/pH6.6) with different internal controls. **m.** Binding profile of StpA at *proVWX* operon and its flanking genes at pH 6.6 in the presence and absence of H-NS strains. **n.** Fold change of H-NS relative binding efficiency (Δ*hns*/WT) with different internal controls. The binding efficiency of H-NS/StpA at *hdeD*1, *dnaA* and *appB* were used for data normalization. The presented data are using *hdeD*1 as the reference. Statistical significance was determined by unpaired, two-tailed Student’s *t*-test. *p*-values indicate the level of significance: *p* < 0.05 (\**), p < 0.01 (**), p < 0.001 (\*\*\**) and *ns* indicates no significant difference.

To further investigate the pH-insensitivity of *proVWX*, we performed ChIP-qPCR to map the binding profiles of H-NS and StpA across the *proVWX* operon and its flanking regions. H-NS and StpA exhibited the strongest binding signals at the promoter region, consistent with previous ChIP-chip results (23). Notably, in addition to the major binding peaks of H-NS at its high-affinity sites, a minor H-NS binding peak was also detected downstream at the flanking gene *ygaY* **(Figure 5g, i and supplementary figure 9a)**, a site previously reported in some other studies (23), but not consistently present in all (112). The appearance of this weak binding signal supports the previously proposed model in which H-NS mediates DNA looping of *proVWX* via an interaction with the *ygaY* region (18). In contrast, this minor peak was absent in the StpA binding profile **(Figure 5g, i)**, suggesting that this site represents an H-NS-specific binding locus. H-NS binding to the *proVWX* region exhibited a pH-responsive decrease in signal upon acid shock **(Figure 5g, h, supplementary figure 9a, b)**. Deletion of *stpA* did not affect the DNA-binding capacity of H-NS **(Figure 5i, j)**. Interestingly, the binding signal of StpA at *proVWX* operon does not appear to be affected by acid shock **(Figure 5k, l, supplementary figure 9c, d)**. Upon deletion of *hns*, StpA binding was markedly enhanced across the *proVWX* regulatory region and its flanking sequences **(Figure 5m, n)**, suggesting that StpA may partially compensate for the absence of H-NS in regulating this locus.

Finally, we mapped the local chromatin conformation of the *proVWX* operon by using 3C-qPCR and analysis of published Micro-C data. To facilitate comparison with previous 3C-qPCR studies (18), we used the same restriction enzyme (NIaIII) to digest the operon, as well as the same primer pairs and TaqMan probe for amplification. Data were normalized using the same internal control (Fragment 6), allowing us to compare the relative interaction frequencies between different target fragments and the anchor fragment **(Fragment 3, highlighted in red in figure 6a-e)**. In our earlier work, we identified a prominent hairpin structure at the *proVWX* promoter region, along with a loop formed between a high-affinity H-NS binding site (Fragment 3) and the downstream *ygaY* gene (Fragment 11). We further established a correlation between this chromatin architecture and *proVWX* transcriptional regulation (18).

In the current 3C-qPCR study our earlier observations were reproduced (**Figure 6a**) and the interpretation of these data was strengthened by examination of Micro-C contact maps revealing the presence of a promoter-proximal hairpin-like structure (**Supplementary figure 10a**). However, long-range loop interactions were difficult to discern in the Micro-C contact map (**Supplementary figure 10**). Interestingly, the hairpin-like structure and loop conformation both were not affected by pH shock in the WT strain **(Figure 6a)**. This observation is consistent with the minimal transcriptional changes of *proVWX* under the same conditions **(Figure 5a)**. In the Δ*stpA* strain, however, we observed a localized opening of the chromatin conformation surrounding the promoter region of *proVWX* upon acid shock **(Supplementary figure 11a)**. Unexpectedly, this did not correspond to an increase in transcription; instead, a decrease in transcript levels was detected at *proV* and *proW* **(Supplementary figure 8a)**. We speculate that this may be due to concurrently increased interactions between the anchor fragment and upstream promoter-adjacent regions (fragment 13, 16, 17) **(Supplementary figure 11a)**, which could continue to hinder RNA polymerase access despite the loosening of downstream loops. Similar to our findings with the *hdeAB* operon, the global chromatin architecture of the *proVWX* locus appeared unaffected by acid shock in the Δ*hns* strain **(Supplementary figure 11b)**. Comparative analysis of 3D chromatin structure across different strains revealed that deletion of *stpA* did not disrupt the structural integrity of this operon **(Figure 6b, supplementary 10b)**. With the deletion of *hns*, a significant decrease in the interaction frequency between the two sites forming the loop structure was observed (fragment 3 and 11), indicating a loss of the loop conformation (**Figure 6c**). In the previous study (18), when point mutations were introduced into the high-affinity H-NS binding site (downstream regulatory element (DRE), fragment 3) to reduce H-NS binding affinity, a reduction in the relative interaction frequency between the high-affinity site and the adjacent target region (fragment 2 and 4) was detected. However, this phenomenon was not observed in the *hns* mutant strain, which may be attributed to compensatory binding of StpA to the promoter region in the absence of H-NS (**Figure 5m**). The Micro-C contact maps from the Δ*hns* strains revealed the presence of a residual, small hairpin structure at the promoter region (**Supplementary 10c, black arrow)**. In contrast, this structure was completely abolished in the Δ*hns* Δ*stpA* mutant strains (**Supplementary 10d, black arrow)**, where the chromatin organization more closely resembled an operon-sized chromosomal interaction domain (OPCID) associated with high transcriptional activity (**Supplementary 10d)**. However, in the Δ*hns* Δ*stpA* mutant as well as in the Δ*hns* Δ*stpA* Δ*rpoS* triple mutant strains, 3C-qPCR analysis detected only a pronounced loss of long-range loop interactions, without changes in promoter-proximal structure **(Figure 6d-e)**. This discrepancy may be attributable to the relatively large size of fragment 2 used in the 3C-qPCR assay, which lacks the resolution necessary to capture subtle structural alterations when compared with the 50 bp resolution of Micro-C contact maps. Moreover, Micro-C data revealed frequent interactions between a high-affinity H-NS binding site and the terminator region of the upstream *nrdF* gene, a region encompassed within fragment 2, which may have further obscured detection of local promoter-associated structural changes in the 3C-qPCR assay.

**Figure 6.**
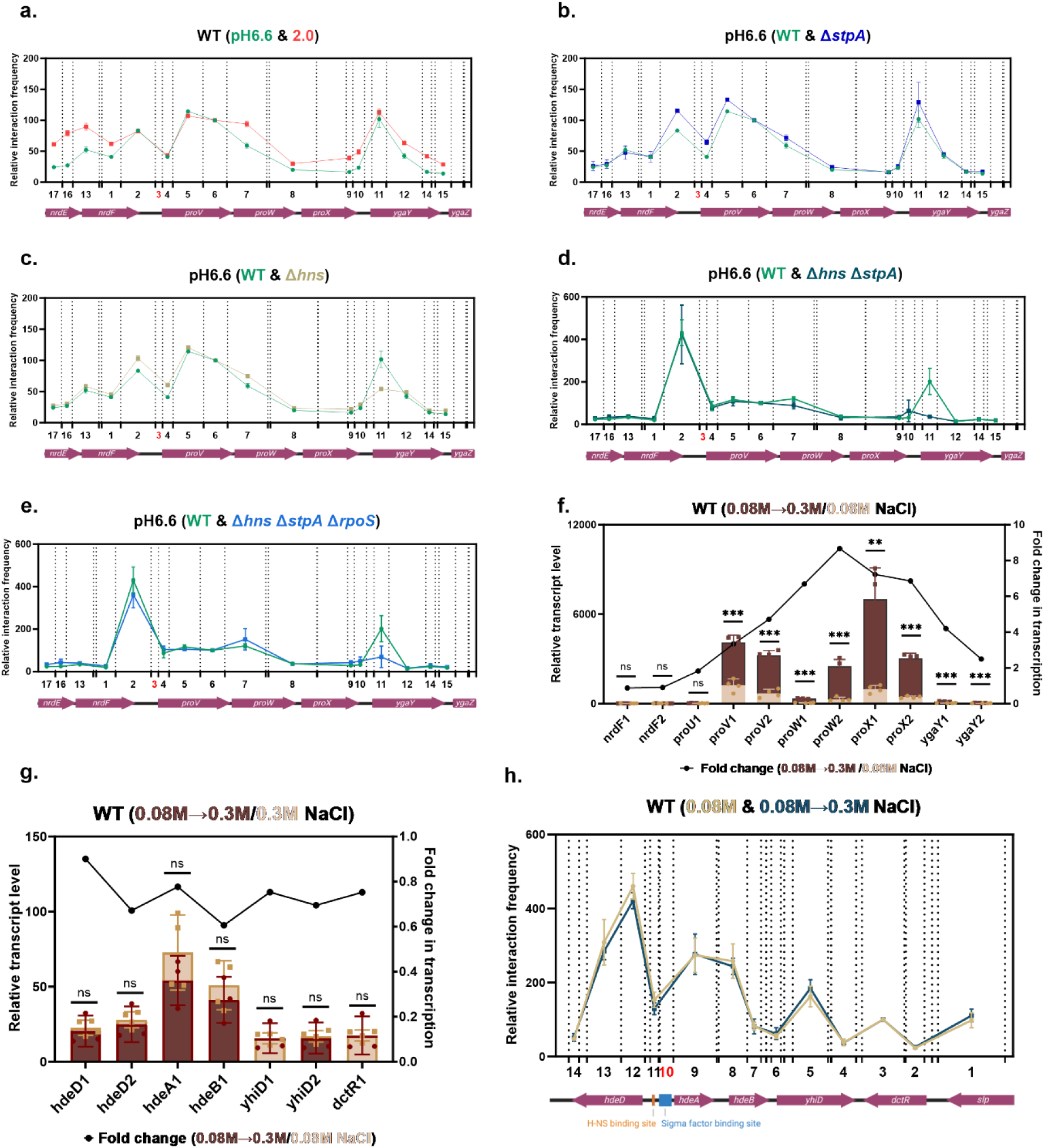
Differential responses of the *proVWX* and *hdeAB* operons to shock condition. **a.** Relative interaction frequency at *proVWX* operon and flanking genes in the WT strain at pH6.6 and pH2.0 conditions. Relative interaction frequency at the *proVWX* operon and flanking genes at pH6.6 condition in Δ*stpA* (**b**), Δ*hns* (**c**), Δ*hns* Δ*stpA* (**d**) and Δ*hns* Δ*stpA* Δ*rpoS* (**e**) strains compared with the WT. Dashed lines indicate NlaIII restriction sites within the *proVWX* operon. The x-axis shows the genomic positions and lengths of tested target fragments and the anchor fragment (fragment 3, marked in red). The y-axis represents relative interaction frequencies between target fragments and the anchor. The interaction frequency between anchor fragment and fragment 6 was used for data normalization. Transcription profile of the *proVWX* operon (f), *hdeAB* operon (g) and its flanking genes at low -salt (0.08 M NaCl) and salt shock (0.08 M NaCl →to 0.3 M NaCl) conditions. The binding efficiency of H-NS/StpA at *hdeD*1, *dnaA* and *appB* were used for data normalization. The presented data are using *hdeD*1 as the reference. Statistical significance was determined by unpaired, two-tailed Student’s *t*-test. *p*-values indicate the level of significance: *p* < 0.05 (\**), p < 0.01 (**), p < 0.001 (\*\*\**) and *ns* indicates no significant difference. Relative interaction frequency of *hdeAB* operon and its flanking genes at low salt and salt shock conditions in WT strain (h). Dashed lines indicate the MluCI restriction enzyme cleavage sites within the *hdeAB* operon. The x-axis shows the relative positions and lengths of target fragments and the anchor fragment (fragment 10, highlighted in red) used in this study. The y-axis represents the relative interaction frequency between the anchor and target fragments.

Taken together, these findings provide further evidence for the essential role of H-NS in the formation of the chromatin hairpin-like and loop structure at the *proVWX* operon. Additionally, StpA can function to maintain local chromatin structure and repress transcription in the absence of H-NS or when H-NS binding at the *proVWX* promoter region is reduced. At the same time, Micro-C contact maps revealed the emergence of long-range loop interactions at the *proVWX* operon upon deletion of *hns* or simultaneous deletion of *hns* and *stpA* **(Supplementary 10c-d)**. These observations highlight the complexity of chromatin architecture and transcriptional regulation in the vicinity of *proVWX* and suggest that additional nucleoid-associated proteins may be involved in its regulation.

### Salt shock does not disrupt the H-NS–maintained chromatin architecture at the *hdeAB* locus

To further investigate the conservation of this chromatin structure-mediated transcription regulation mechanism in the context of *E. coli*’s specific transcriptional responses to environmental signals, we mapped the changes in transcription landscape and local three-dimensional chromatin structure of the *hdeAB* operon following salt shock (10 minutes shock from 0.08 M NaCl to 0.3 M NaCl). To ensure that the salt shock was sufficient to trigger a transcriptional response in *E. coli*, we first examined the expression of the osmotically responsive *proVWX* operon as a control. Consistent with our previous results (18), salt shock induced a strong upregulation of *proVWX* transcription (3–8 fold) **(Figure 6f)**. While transcription of the upstream gene *nrdF* was not affected by osmotic stress, the downstream gene *ygaY* exhibited a moderate increase **(Figure 6f)**, likely due to transcriptional read-through from the *proVWX* operon by RNA polymerase. In the Δ*rpoS* single mutant, 7-14 fold upregulation of *proVWX* transcription was observed **(Supplementary figure 12a)**. However, in the Δ*hns* Δ*stpA* and Δ*hns* Δ*stpA* Δ*rpoS* strains, the transcriptional response of *proVWX* to salt shock was markedly reduced, exhibiting no or only a mild (1–3 fold) increase **(Supplementary figure 12b, c)**. This resembles the loss of the stress-response in the Δ*hns* Δ*stpA* and Δ*hns* Δ*stpA* Δ*rpoS* mutants similar to the non-pH-responsive behavior of *hdeAB* transcription observed in the same mutant strains. It can be explained in the same way: the transcription repression of *proVWX* from H-NS and StpA is relieved in their absence and the slight increase in transcription of *proVWX* is attributed to the inherent osmosensitivity of *proVWX* (95). Compared with the WT under low salt conditions (0.08 M NaCl), deletion of *rpoS* had no effect on the transcription of *proW*, *proX*, or the flanking genes, whereas the transcription of *proV* was significantly reduced **(Supplementary figure 12d)**. Under the same conditions, only a subset of amplicons in the Δ*hns* Δ*stpA* and Δ*hns* Δ*stpA* Δ*rpoS* strains exhibited mild transcriptional upregulation **(Supplementary figure 12e, f**) compared with the WT strain, which is generally consistent with the results we described earlier (**Figure 5d, e**). Taken together, these results demonstrate that the transcription induction of *proVWX* is osmo-stress specific and insensitive to acid shock.

Next, we examined the level of transcription across and flanking the *hdeAB* operon. The results showed that *hdeAB* transcription was not activated by salt shock in any of the tested strains **(Figure 6g and supplementary figure 13a-e)**. Under low-salt conditions (0.08 M NaCl), compared with the WT strain, the transcription of *hdeAB* was decreased in the Δ*rpoS* mutant (**Supplementary figure 13f**), the transcription of *hdeA* and *hdeB* was increased approximately 1.5-fold in the Δ*stpA* mutant, with no significant changes observed in the flanking genes (**Supplementary figure 12g**). In the Δ*hns*, Δ*hns* Δ*stpA* and Δ*hns* Δ*stpA* Δ*rpoS* strains, the transcription levels of all examined genes were elevated by approximately 15- to 75-fold relative to the WT (**Supplementary figure 13h-j**). In the acid shock series experiment, we established a correlation between the transcriptional activity of the *hdeAB* operon and its hairpin structure at the promoter region **(Figure 1, 2, 3, 4 and supplementary figure 5)**. We also examined the three-dimensional chromatin organization of the *hdeAB* operon under different osmotic conditions in the WT, Δ*stpA* and Δ*hns* strains. As expected, the hairpin-like chromatin structure of *hdeAB* operon could still be observed **(Figure 6h)**, but the structure was not altered following salt shock in all strains tested **(Figure 6h and supplementary figure 14a, b)**. Moreover, consistent with our observations under pH 6.6 conditions, the hairpin-like structure at the *hdeAB* promoter region was maintained even in the absence of StpA **(Supplementary figure 14c)**, whereas deletion of *hns* resulted in a transition from the hairpin-like structure to a more open conformation at low salt conditions **(Supplementary figure 14d)**. These results demonstrate that the hairpin-like structure maintained by H-NS within the *hdeAB* operon and its flanking regions does not undergo a transition to an open conformation upon salt shock, unlike what is observed under acid stress. This further underlines the essential role of the H-NS maintained hairpin structure within the *hdeAB* operon in regulating its transcription.

## Discussion

The bacterial nucleoid is organized and compacted by nucleoid-associated proteins, DNA topology, and macromolecular crowding (12). During biological processes such as transcription, DNA replication and chromosome segregation, the nucleoid undergoes continuous structural changes (113–116). As a prototypical nucleoid-associated protein, H-NS has been shown to play dual roles in maintaining chromatin architecture and regulating transcription (6, 12, 36). In this study, by integrating *in vivo* and *in vitro* approaches, we demonstrate that H-NS functions as a direct sensor of acidic environments and regulates transcription of the acid-resistance operon *hdeAB* by remodeling a promoter-proximal hairpin structure, thereby enhancing the survival of *E. coli* under acidic conditions **(Figure 7a)**. This is analogous to our previous model in which H-NS regulates transcription of the osmo-regulated *proVWX* operon by local three-dimensional remodeling of chromatin architecture (18), establishing this mechanism as potentially generic for H-NS mediated responses to physico-chemical changes. **(Figure 7a-b)**. Moreover, in light of *in vitro* evidence demonstrating that H-NS responds to diverse environmental signals, we propose that chromatin restructuring in response to environmental cues is conserved at H-NS-repressed target genes. We expect that similar regulatory mechanisms operate in other bacterial species that harbor H-NS or H-NS-like proteins.

**Figure 7:**
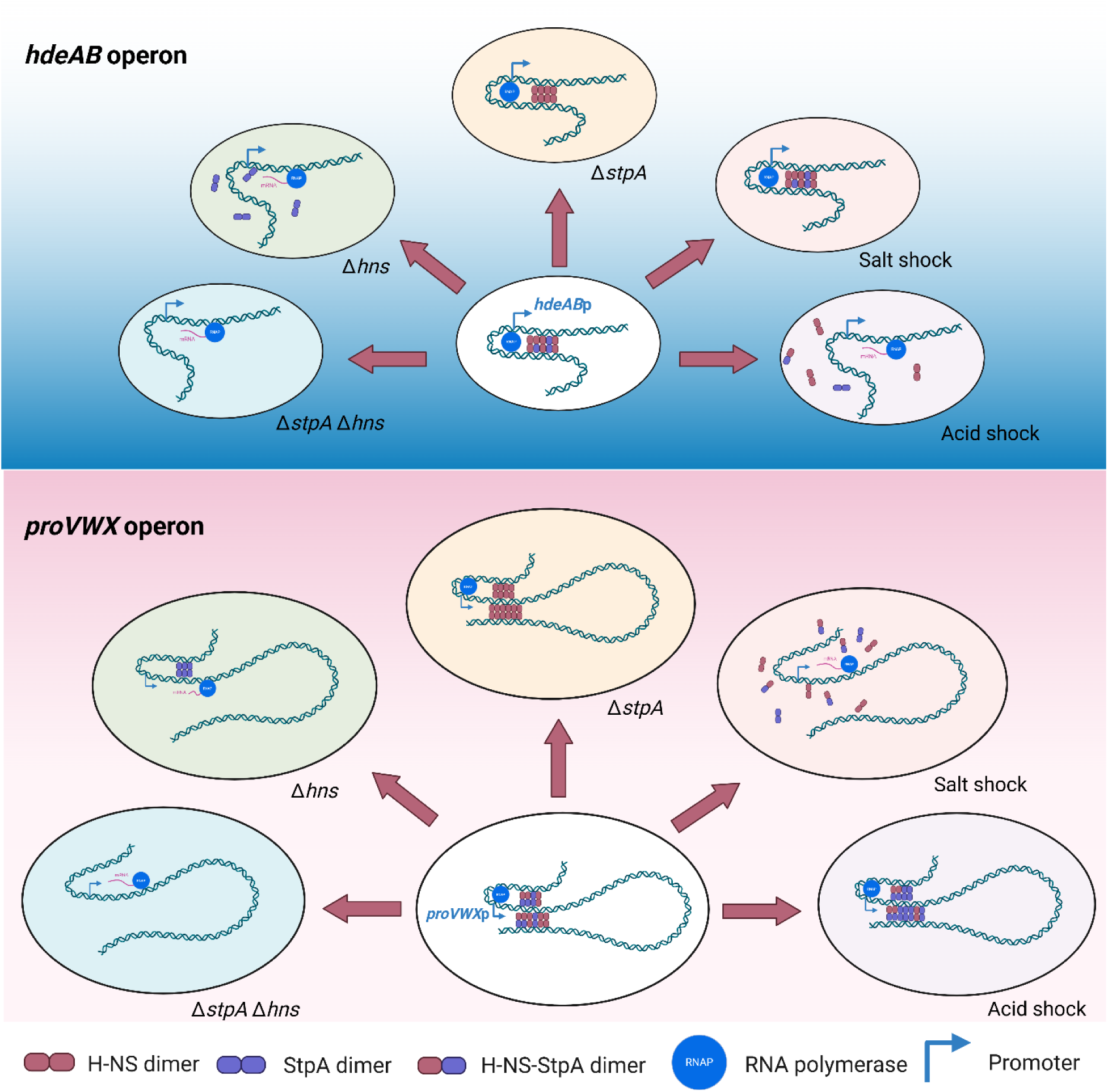
Models of regulation of the *hdeAB* and *proVWX* operon. Local chromatin architecture of the *hdeAB* and *proVWX* operons under different environmental conditions and in different genetic backgrounds determines operon transcription.

In addition, the unique promoter-proximal hairpin-like structure both identified in our 3C-qPCR analyses **(Figure 3a and 6a)** and in published Micro-C data *in vivo* (109)(**Supplementary figure 6 and 10**), extends our understanding of the role of H-NS and StpA in bacterial chromatin organization. In this study, the hairpin-like structure formed at the *hdeAB* promoter region is primarily dependent on H-NS, as deletion of *stpA* alone does not disrupt this architecture **(Figure 3e, Supplementary figure 6)**. Loss of H-NS or acid-induced reduction of H-NS binding leads to loss of the hairpin-like structure, resulting in a more ‘open’ chromatin conformation at the promoter **(Figure 3a, f and Supplementary figure 6)**. This ‘open’ conformation increases promoter access for RNA polymerase, thereby enhancing transcription. The chromatin architecture of the *proVWX* region is considerably more complex. In *hns* mutant or *hns stpA* double mutant strains, the loop formed in the WT strain between the high-affinity H-NS binding site and the downstream *ygaY* gene is not present (18) **(Figure 6c, d)**, accompanied by the emergence of increased long-range chromosomal interactions (**Supplementary figure 10**). In another recent study, it was shown using Hi-C that deletion of *stpA* increases H-NS-mediated bridging interactions(117), similar to the trend observed in our own experiments at the *proVWX* operon **(Figure 6b)** but not at the *hdeAB* operon. Furthermore, during the stationary phase, deletion of *fis* or *ihf* also enhances H-NS-mediated DNA bridging (117). This indicates that a more complex network of physical interactions or structural interplay between NAPs is involved in maintaining chromatin architecture.

With respect to StpA, our model indicates that StpA can cooperate with H-NS to repress the transcription of both the *hdeAB* and *proVWX* operons; however, its repressive capacity is clearly weaker than that of H-NS **(Figure 1g, h and Figure 5b, c)**. The relatively weak repressive effect of StpA on *hdeAB* transcription is attributed to its low cellular abundance, as H-NS strongly suppresses *stpA* transcription **(Supplementary figure 4a)**. In addition, in the absence of H-NS, StpA alone is insufficient to maintain the hairpin-like higher-order chromatin structure at the *hdeAB* locus, whereas it can partially preserve the hairpin structure at the *proVWX* region (**Supplementary figure 6 and 10)**. These observations suggest that when StpA cannot form heterodimers with H-NS, it may exhibit more refined binding preferences among different genomic loci, even within AT-rich regions. This interpretation is readily supported by previous ChIP data (23). Accordingly, under H-NS-deficient conditions, StpA remains capable of independently maintaining hairpin-like chromatin structures at genomic sites where it displays high binding affinity (e.g. *yfjW*, *ygeH*, *proVWX*, *slp-dctR*, *yhiS*) (23, 109) **(Figure 2g and Supplementary figure 6)**. However, the molecular basis underlying StpA’s binding preference, within AT-rich genomic regions, remains to be elucidated. Over the past five years, evidence has been increasing that post-translational modifications (PTMs) of H-NS can directly influence the transcription of its target genes (29) and ChIP-seq analyses have mapped genome-wide H-NS binding profiles under different PTM states (22, 37). However, evidence directly linking the chromatin structures maintained by H-NS and StpA with their transcription regulatory functions remains limited.

Another important question that we addressed is whether H-NS, as a global regulator of many environment-responsive genes (29–33), selectively modulates transcription of specific target operons in response to different environmental cues and if so, by what mechanism. To examine this, we measured transcriptional responses of the osmotic-stress-responsive *proVWX* operon and assessed the local chromatin architecture of the *proVWX* locus under different pH conditions. We found that neither the transcription level of *proVWX* nor its local chromatin conformation was altered by pH shock. However, ChIP-qPCR revealed a marked reduction in H-NS binding at the *proVWX* promoter region following acid treatment (**Figure 5g, h, supplementary figure 9a, b**). We attribute the maintenance of *proVWX* transcription and local chromatin structure under acid stress to compensatory binding by StpA at the promoter, an idea supported by the pronounced increase in StpA occupancy at the promoter in the Δ*hns* strain (**Figure 5m, n)**. The compensatory binding of StpA under acidic conditions is consistent with its low sensitivity to pH changes in DNA binding (65). In the *hns stpA* double mutant strain, the osmotic-stress-induced transcription of the *proVWX* operon is driven by the intrinsic osmosensitive properties of the *proVWX* promoter (95, 118, 119), thereby conferring specificity in the transcriptional response to environmental cues. Notably, we did not observe an increase in StpA binding at the *hdeAB* promoter in the Δ*hns* strain (**Figure 2g, h)**, which can be explained by the DNA-binding preferences of StpA described above (23). Together, these findings support a model in which precise, stimulus-specific transcriptional regulation of the *proVWX* operon is achieved through differential sensitivities of H-NS and StpA to acidic signals, coupled with compensatory binding of StpA at the *proVWX* promoter region. The transcription of the *hdeAB* operon exhibits a specific response to environmental cues, as well. Specifically, following salt shock, neither its transcriptional activity nor its local chromatin architecture exhibits detectable changes (**Figure 6g, h)**.

In summary, our study demonstrates that the transcription of stress-responsive genes exhibits specific responses to distinct environmental cues and provides evidence of a key role for the global regulators H-NS and StpA in coordinately modulating higher-order chromatin architecture to control gene transcription. In addition, our findings expand the current understanding of the capacity of StpA to exert regulatory functions independent of H-NS *in vivo*.

## Data availability

All data generated in this study have been deposited in the 4TU Repository (Doi: 10.4121/f75943dc-4404-4784-8445-ceee37f141ad). The RT-qPCR, ChIP-qPCR and 3C-qPCR data generated in this study are also provided in the Supplementary files. The Micro-C data are available under the Gene expression Omnibus accession code GES272161.

## Supplementary Data

Supplementary Data are available at NAR Online.

## Acknowledgments

We thank Ilya Shamovsky and Evgeny Nudler for providing the Micro-C data for supplementary figure 6, 10. We thank all the students, Nick Fang, Tijn Kops, and Xuan Zhang, for their contributions to this study during their internships. We thank Anneloes Cramer-Blok and Monika Timmer, laboratory technician, for their daily support during this study. Graphical abstract was created in BioRender. Ge, P. (2026) https://BioRender.com/5qabkxs.

## Author Contributions

Conceptualization: P.G., F.M.R., R.T.D.; Formal Analysis: P.G., M.T., R.T.D.; Funding Acquisition: R.T.D., P.G., B.P.; Investigation: P.G., F.M.R., L.K.F.G., M.K.M.C., M.T.; Methodology: P.G., F.M.R., R.T.D.; Project Administration: R.T.D.; Resources: P.G., F.M.R., K.S.; Software: M.T.; Supervision: R.T.D., B.P., K.S.; Validation: P.G., L.K.F.G., M.K.M.C., M.T.; Visualization: P.G., M.T.; Writing – Original Draft Preparation: P.G., F.M.R., R.T.D.; Writing – Review & Editing: All of authors.

## Funding

This work was supported by the Dutch Research Council [OCENW.GROOT.2019.012 to RTD], China Scholarship Council [grant no. 202103250002 to PG] and European Research Council [ERC Starting Grant 950655-Silencer to MT and BP].

## Conflict of interest disclosure

None declared.

## Notes

### Competing Interest Statement

The authors have declared no competing interest.

