## Supplementary Files for "Adaptive gene transcription in *Escherichia coli* under environmental stress"

#### Supplementary figures

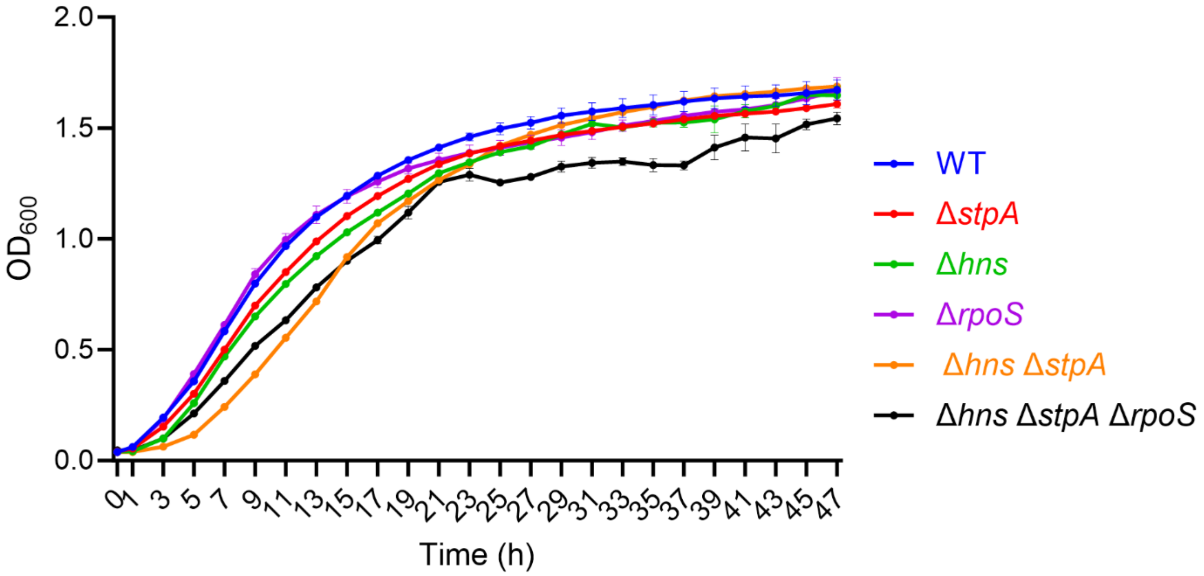


**Supplementary figure 1:** The growth curve of strains used in this study. All of strains were cultured in M9 medium (pH 6.6). Data are presented as Mean ± SD. OD600, optical density at 600 nm.

Classified as Internal | Intern

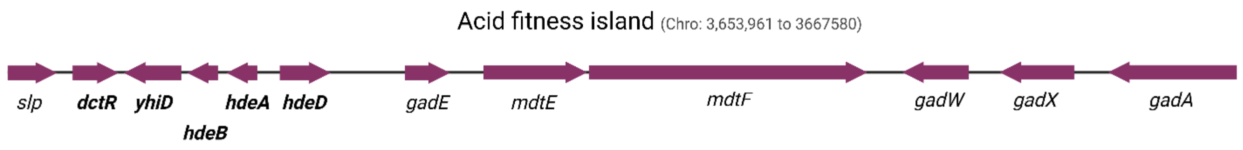


**Supplementary figure 2:** Schematic showing the map of the acid fitness island genes in *E. coli* MG1655 (NC_000913.3).

Classified as Internal | Intern

a.

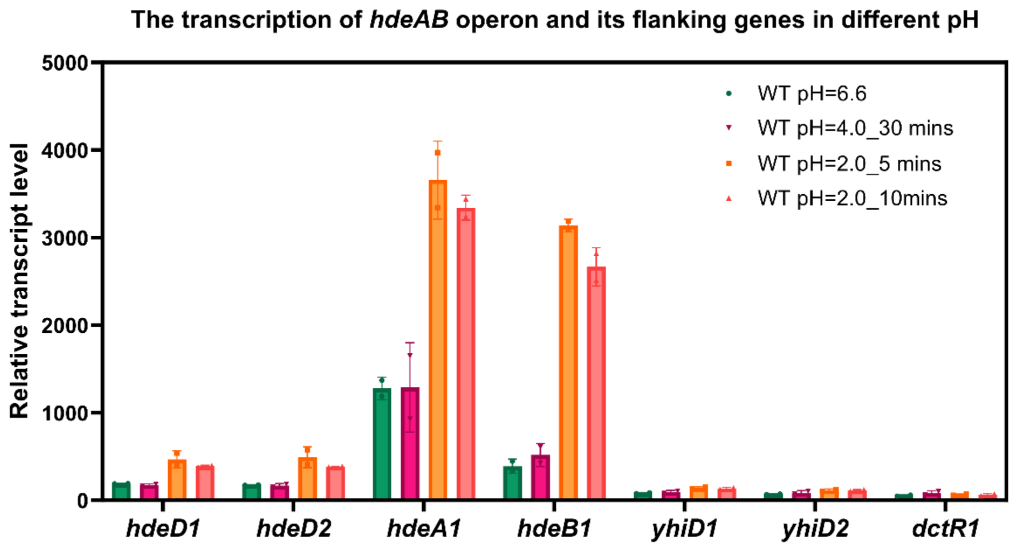


b.

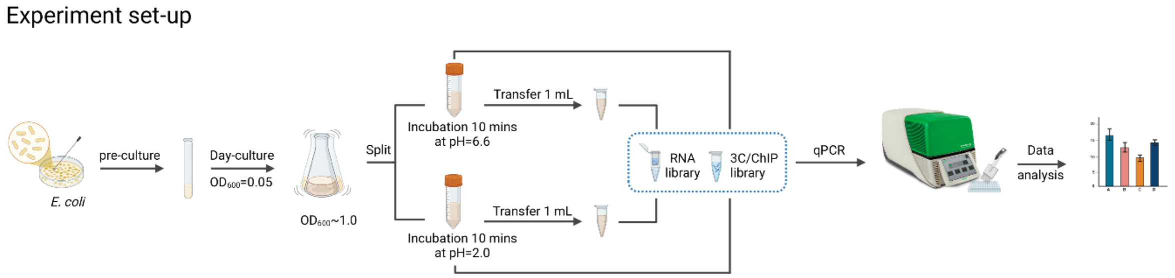


**Supplementary figure 3:** Transcription profile of the *hdeAB* operon and its flanking genes with different acid shock time in WT strain **(a)**. Experiment set-up for RT-qPCR, ChIP-qPCR and 3C-qPCR **(b)**.

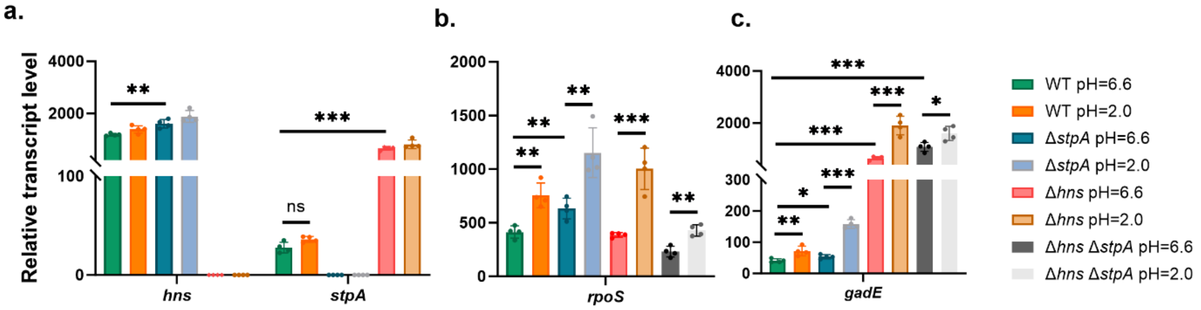


**Supplementary figure 4:** Transcription profile of the *hns* (**a**), *stpA* (**a**) *rpoS* (**b**) and *gadE* (**c**) at pH 6.6 and pH 2.0 conditions in the WT, Δ*stpA*, Δ*hns* and Δ*hns* Δ*stpA* strains. Each group includes four biological replicates and three technical replicates. *DnaA* is used for data presentation. Statistical significance was determined by unpaired, two-tailed Student's t-test. p-values indicate the level of significance: $p < 0.05$ (*), $p < 0.01$ (**), $p < 0.001$ (***) and *ns* indicates no significant difference.

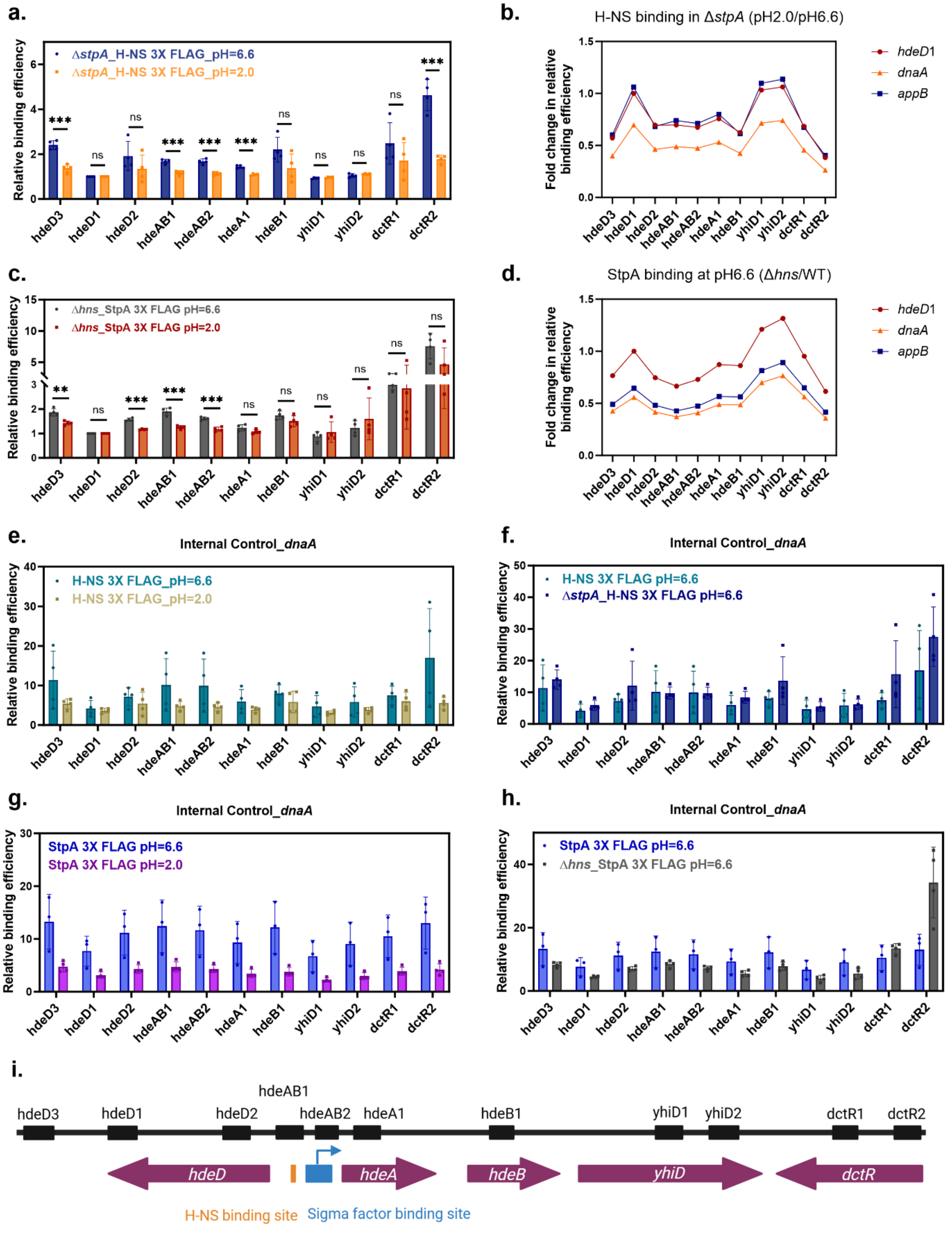


**Supplementary figure 5**: **a.** Binding profile of H-NS at *hdeAB* operon and its flanking genes at pH 6.6 and pH 2.0 conditions in Δ*stpA* strain. **b.** Fold change of H-NS relative binding efficiency (pH2.0/pH6.6) with different

Classified as Internal | Intern

internal controls. **c.** Binding profile of StpA at *hdeAB* operon and its flanking genes at pH 6.6 and pH 2.0 conditions in Δ*hns* strain. **d.** Fold change of StpA relative binding efficiency (pH2.0/pH6.6) with different internal controls. **e.** Binding profile of H-NS at *hdeAB* operon and its flanking genes at pH 6.6 and pH 2.0 conditions (*dnaA* as internal control). **f.** Binding profile of H-NS at *hdeAB* operon and its flanking genes at pH 6.6 in the presence and absence of StpA (*dnaA* as internal control)**. g.** Binding profile of StpA at *hdeAB* operon and its flanking gene at pH 6.6 and pH 2.0 conditions (*dnaA* as internal control). **h.** Binding profile of StpA at *hdeAB* operon and its flanking gene at pH 6.6 in the presence and absence of H-NS (*dnaA* as internal control). **i.** Schematic showing the positions of amplicons used in ChIP-qPCR analysis. Each group includes four biological replicates with two technical replicates each. Statistical significance was determined by unpaired, two-tailed Student's *t*-test. *p*-values indicate the level of significance: $p < 0.05$ (*), $p < 0.01$ (**), $p < 0.001$ (***) and *ns* indicates no significant difference.

Classified as Internal | Intern

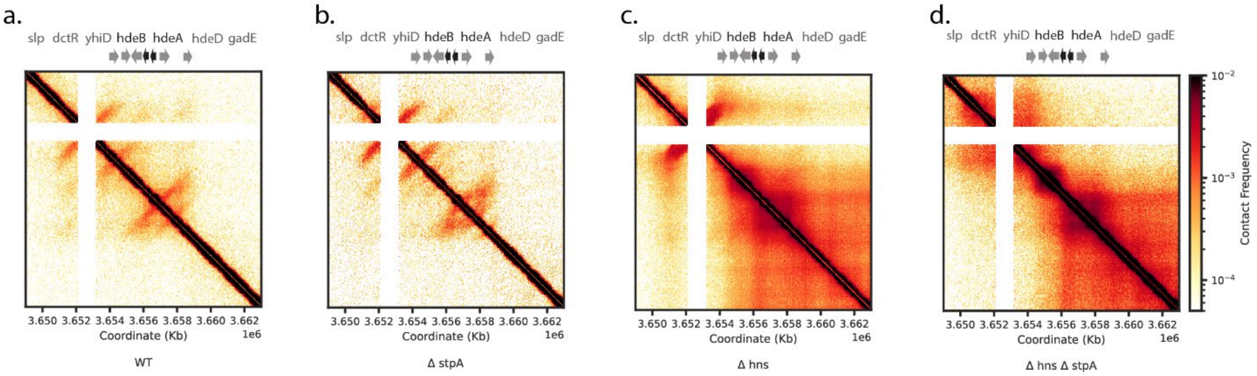


**Supplementary figure 6:** Micro-C contact map of *hdeAB* operon and its flanking genes in WT, Δ*stpA*, Δ*hns* and Δ*hns* Δ*stpA* strains.

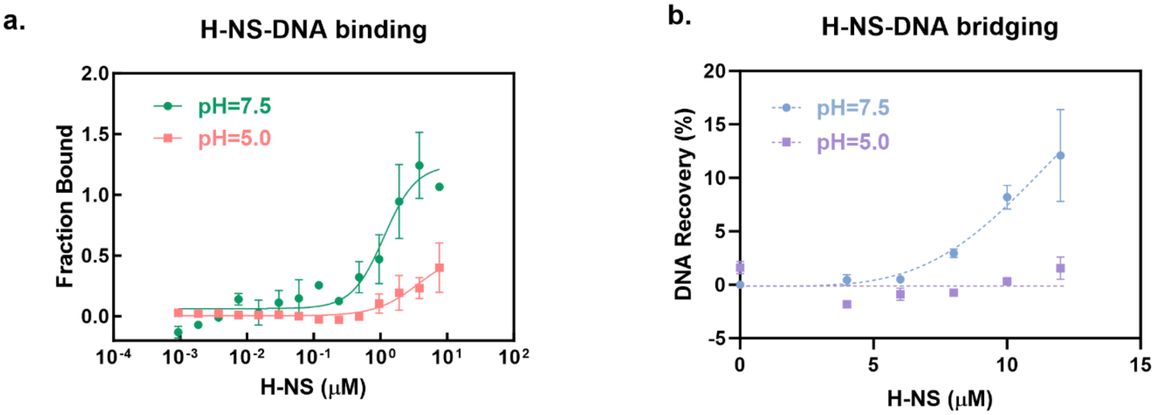


**Supplementary figure 7:** DNA binding of H-NS at pH 7.5 and pH 5.0 (a). For each group with three biological replicates. The DNA bridging efficiency of H-NS at pH 7.5 and pH 5.0. Dashed lines serve as lines to guide the eye (b). For each group with two replicates.

Classified as Internal | Intern

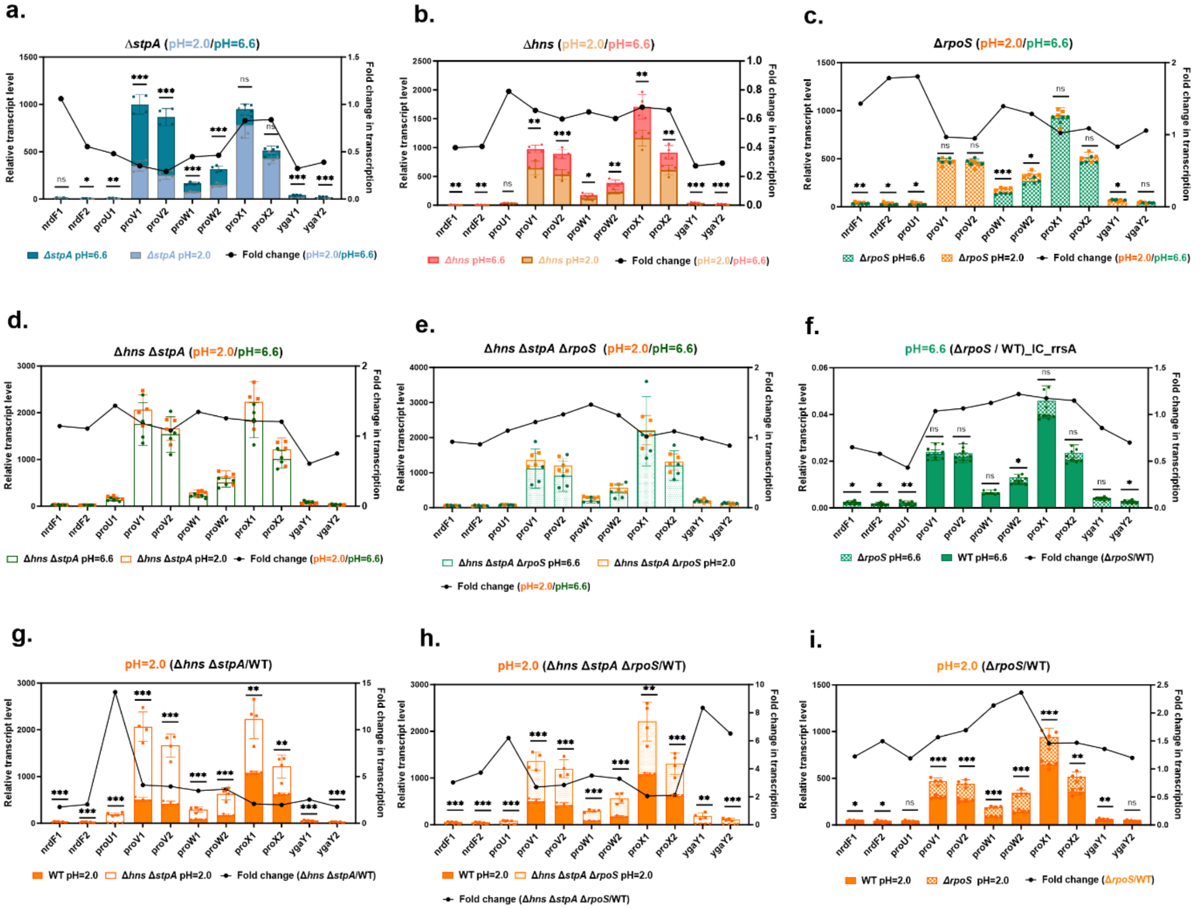


**Supplementary figure 8:** Transcription profile of the *proVWX* operon and its flanking genes at pH 6.6 and pH 2.0 conditions in the Δ*stpA* **(a)**, Δ*hns* **(b)**, Δ*rpoS* **(c)**, Δ*hns* Δ*stpA* **(d)** and Δ*hns* Δ*stpA* Δ*rpoS* **(e)** strains. **f.** Transcription profile of *proVWX* operon and its flanking genes at pH 6.6 in Δ*rpoS* compared with WT (*rrsA* as internal control). Transcription profile of *proVWX* operon and its flanking genes at pH 2.0 in Δ*hns* Δ*stpA* **(g)**, Δ*hns* Δ*stpA* Δ*rpoS* **(h)**, Δ*rpoS* **(i)** compared with WT. Each group includes four biological replicates with three technical replicates each. Statistical significance was determined by unpaired, two-tailed Student's t-test. p-values indicate the level of significance: $p < 0.05$ (*), $p < 0.01$ (**), $p < 0.001$ (***) and *ns* indicates no significant difference.

Classified as Internal | Intern

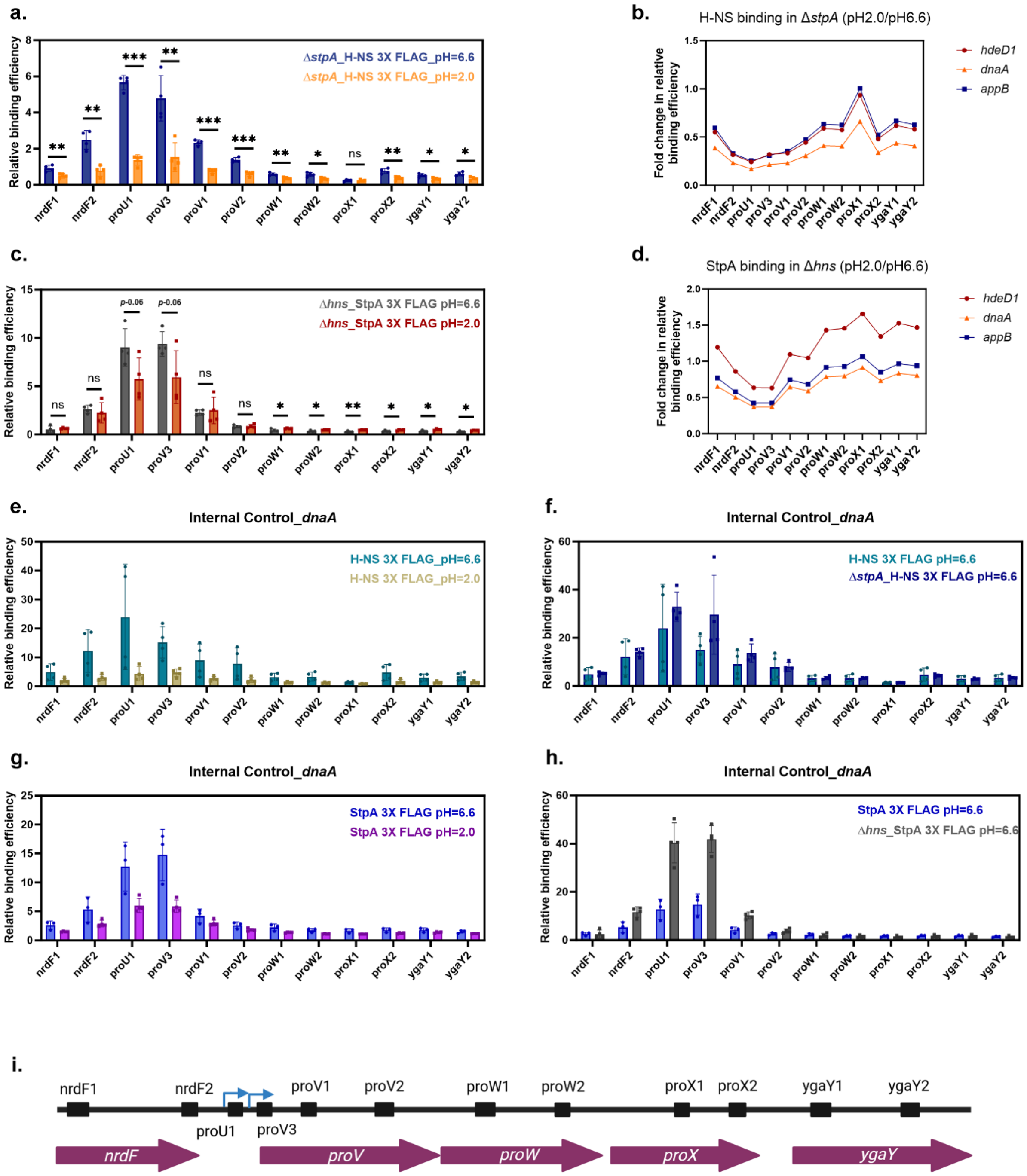


**Supplementary figure 9: a.** Binding profile of H-NS at *proVWX* operon and its flanking genes at pH 6.6 and pH 2.0 conditions in Δ*stpA* strain. **b.** Fold change of H-NS relative binding efficiency (pH2.0/pH6.6) with different internal controls. **c.** Binding profile of StpA at *proVWX* operon and its flanking genes at pH 6.6 and pH 2.0 conditions in Δ*hns* strain. **d.** Fold change of StpA relative binding efficiency (pH2.0/pH6.6) with different internal controls. **e.** Binding profile of H-NS at *proVWX* operon and its flanking genes at pH 6.6 and pH 2.0 conditions (*dnaA* as internal control). **f.** Binding profile of H-NS at *proVWX* operon and its flanking genes at pH

Classified as Internal | Intern

6.6 in the presence and absence of StpA (*dnaA* as internal control). **g.** Binding profile of StpA at *proVWX* operon and its flanking gene at pH 6.6 and pH 2.0 conditions (*dnaA* as internal control). **h.** Binding profile of StpA at *proVWX* operon and its flanking gene at pH 6.6 in the presence and absence of H-NS (*dnaA* as internal control). **i.** Schematic showing the positions of amplicons used in ChIP-qPCR analysis. Each group includes four biological replicates with two technical replicates each. Statistical significance was determined by unpaired, two-tailed Student's *t*-test. *p*-values indicate the level of significance: $p < 0.05$ (*), $p < 0.01$ (**), $p < 0.001$ (***) and *ns* indicates no significant difference.

Classified as Internal | Intern

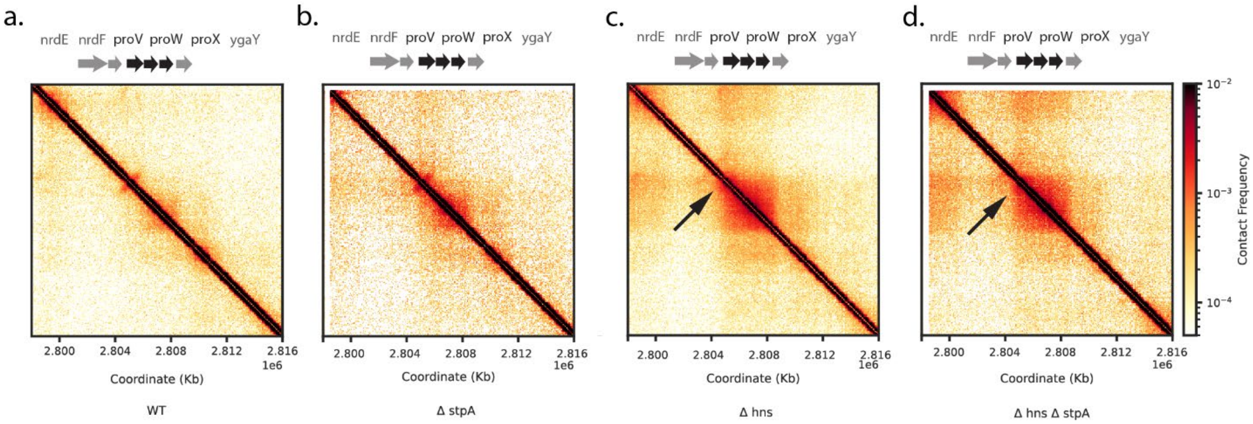


**Supplementary figure 10:** Micro-C contact map of *proVWX* operon and its flanking genes in WT, Δ*stpA*, Δ*hns* and Δ*hns* Δ*stpA* strains.

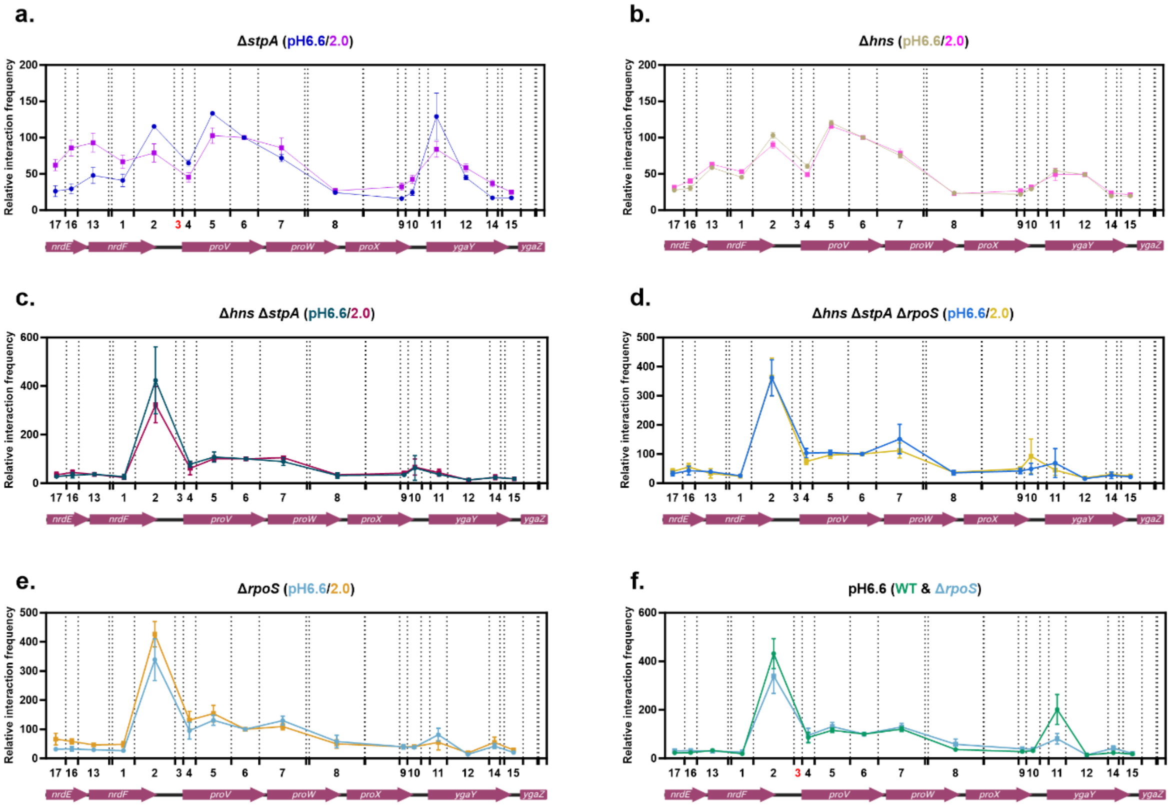


**Supplementary figure 11:** Relative interaction frequency at *proVWX* operon and flanking genes at pH6.6 and pH2.0 conditions in Δ*stpA* **(a)**, Δ*hns* **(b)**, Δ*hns* Δ*stpA* **(c)**, Δ*hns* Δ*stpA* Δ*rpoS* **(d)** and Δ*rpoS* **(e)** strains. Relative interaction frequency at the *proVWX* operon and flanking genes at pH6.6 condition in Δ*rpoS* (**f**)) strain compared with the WT. Dashed lines indicate NlaIII restriction sites within the *proVWX* operon. The x-axis shows the genomic positions and lengths of tested target fragments and the anchor fragment (fragment 3). The y-axis represents relative interaction frequencies between target fragments and the anchor. The interaction frequency between anchor fragment and fragment 6 was used for data normalization.

Classified as Internal | Intern

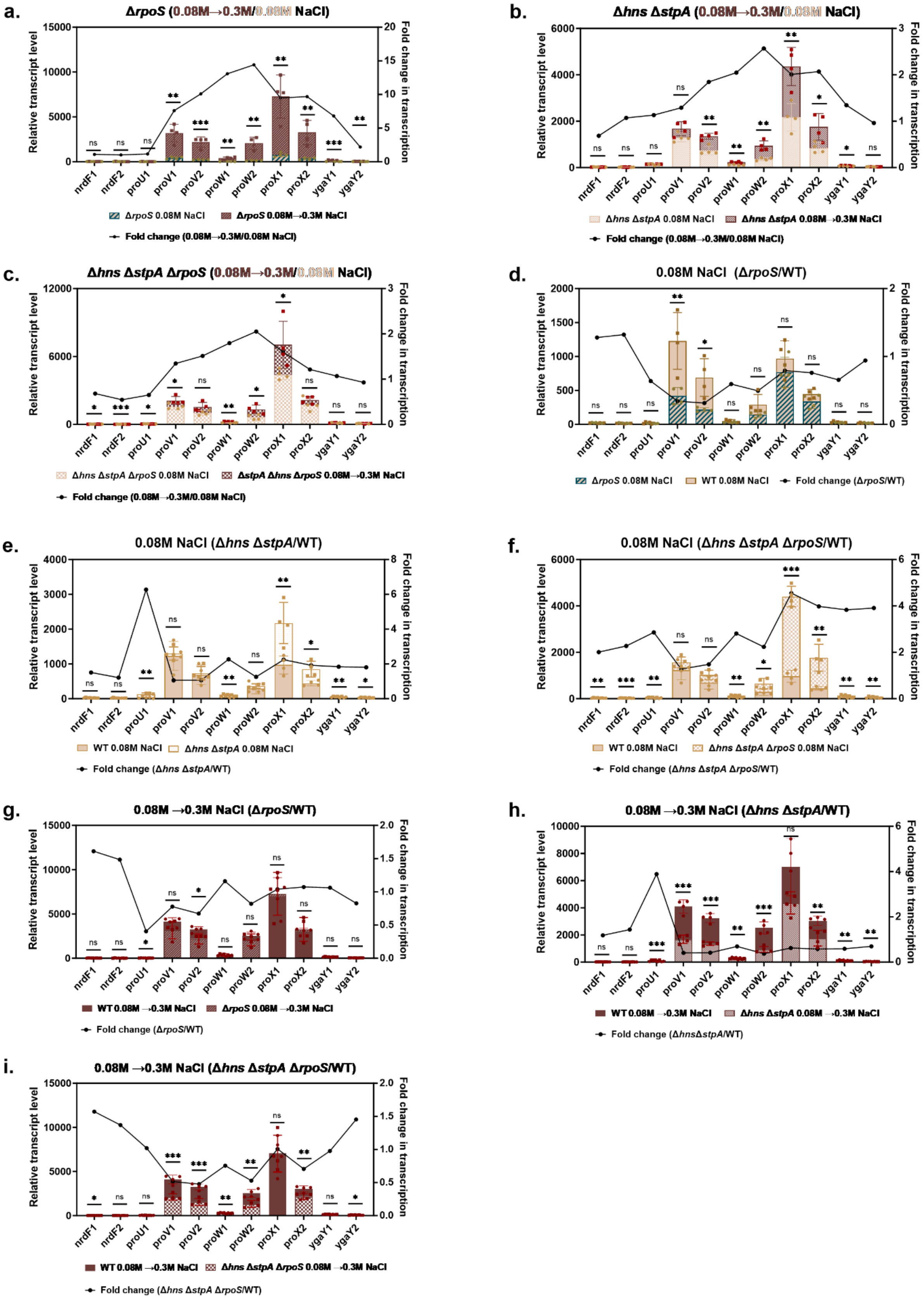


Classified as Internal | Intern

**Supplementary figure 12:** Transcription profile of the *proVWX* operon and its flanking genes at low salt (0.08M NaCl) and salt shock (0.08 M → 0.3 M NaCl) conditions in the Δ*rpoS* (**a**), Δ*hns* Δ*stpA* (**b**) and Δ*hns* Δ*stpA* Δ*rpoS* (**c**) strains. Transcription profile of *proVWX* operon and its flanking genes at low salt (LS) in Δ*rpoS* (**d**), Δ*hns* Δ*stpA* (**e**), Δ*hns* Δ*stpA* Δ*rpoS* (**f**) strains compared with WT. Transcription profile of *proVWX* operon and its flanking genes at salt shock (SS) in Δ*rpoS* (**g**), Δ*hns* Δ*stpA* (**h**), Δ*hns* Δ*stpA* Δ*rpoS* (**i**) strains compared with WT. Each group includes four biological replicates with three technical replicates each. Statistical significance was determined by unpaired, two-tailed Student's *t*-test. *p*-values indicate the level of significance: $p < 0.05$ (*), $p < 0.01$ (**), $p < 0.001$ (***) and *ns* indicates no significant difference.

Classified as Internal | Intern

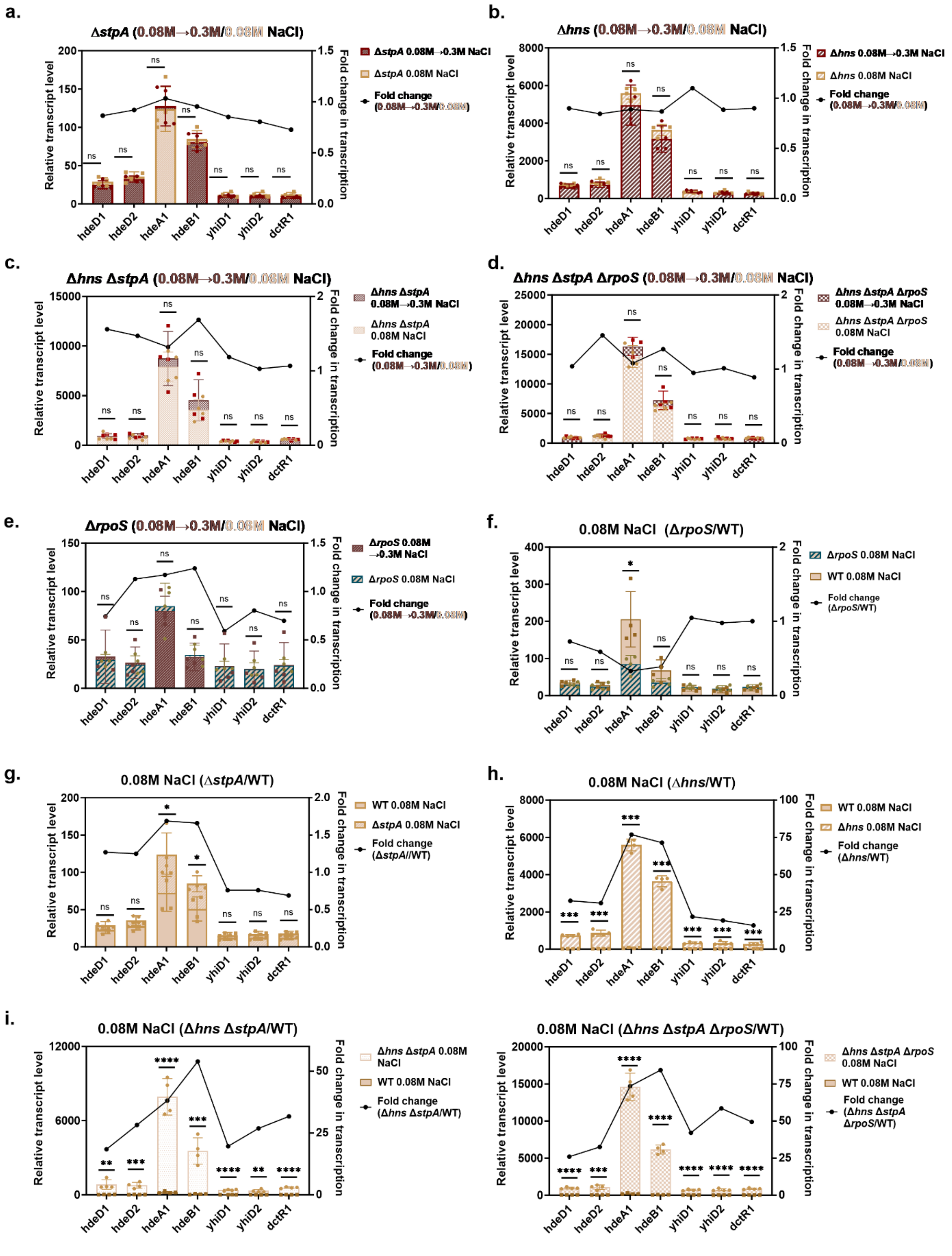


**Supplementary figure 13:** Transcription profile of the *hdeAB* operon and its flanking genes at low salt (0.08M NaCl) and salt shock (0.08M → 0.3M NaCl) conditions in the Δ*stpA* (**a**), Δ*hns* (**b**), Δ*hns* Δ*stpA* (c), Δ*hns* Δ*stpA*

Classified as Internal | Intern

Δ*rpoS* (**d**) and Δ*rpoS* (**e**) strains. Transcription profile of *hdeAB* operon and its flanking genes at low salt (0.08M NaCl) in Δ*rpoS* (**f**), Δ*stpA* (**g**), Δ*hns* (**h**), Δ*hns* Δ*stpA* (**i**) and Δ*hns* Δ*stpA* Δ*rpoS* (**j**) strains compared with WT. Statistical significance was determined by unpaired, two-tailed Student's *t*-test. *p*-values indicate the level of siance: $p < 0.05$ (*), $p < 0.01$ (**), $p < 0.001$ (***) and *ns* indicates no significant difference.

Classified as Internal | Intern

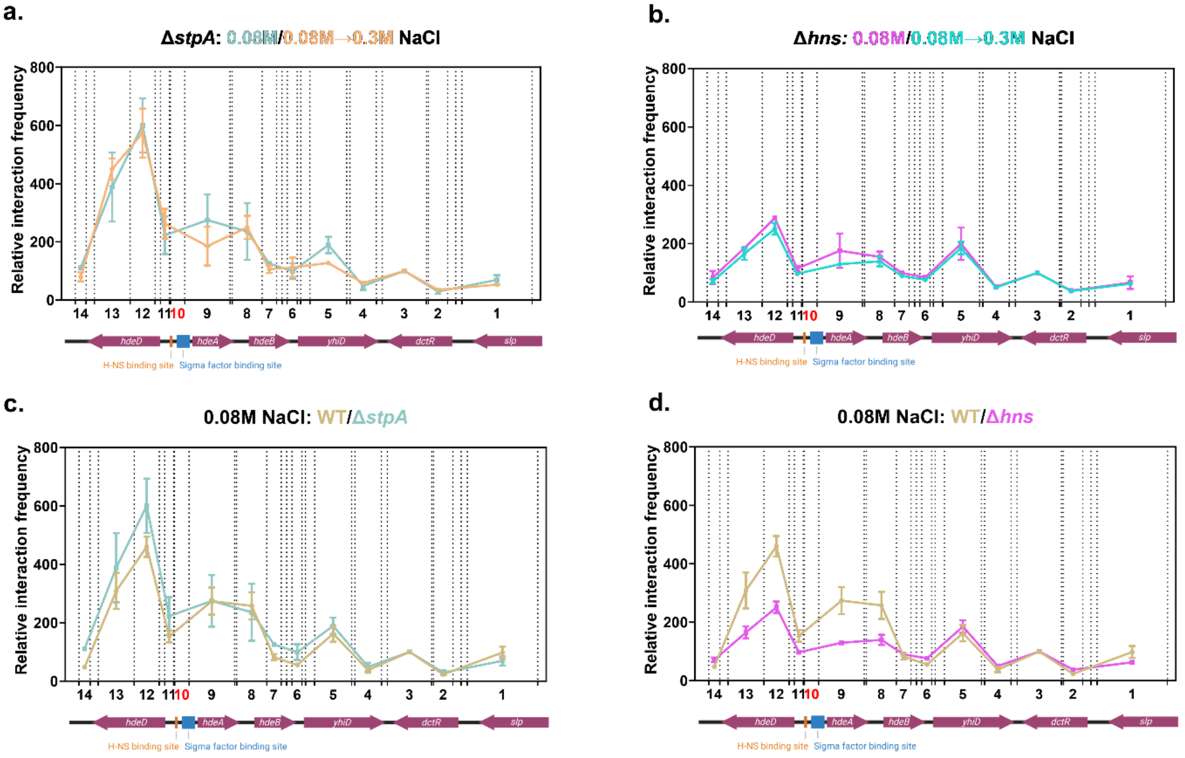


**Supplementary figure 14:** Relative interaction frequency at *hdeAB* operon and flanking genes at low salt (0.08M NaCl) and salt shock (0.08M→ 0.3M NaCl) conditions in Δ*stpA* (**a**) and Δ*hns* (**b**) strains. Relative interaction frequency at the *hdeAB* operon and flanking genes at low salt (0.08M NaCl) condition in Δ*stpA* (**c**) and Δ*hns* (**d**) strains compared with the WT. Dashed lines indicate the MluCI restriction enzyme cleavage sites within the *hdeAB* operon. The x-axis shows the relative positions and lengths of target fragments and the anchor fragment (fragment 10, highlighted in red) used in this study. The y-axis represents the relative interaction frequency between the anchor and target fragments.

Classified as Internal | Intern

### Supplementary tables

#### Table S1. Primers for RT-qPCR

| Primers | Sequences (5'-3') |
|---|---|
| hdeD_RT-qPCR_1.fwd | CCGATGGTGTCTGTAACGC |
| hdeD_RT-qPCR_1.rev | GCTTAACGAACAAACTGGCG |
| hdeD_RT-qPCR_2.fwd | GCAATCCAGTTTATTGCCGTGC |
| hdeD_RT-qPCR_2.rev | AATGCACCCACTACTGTGC |
| hdeA_RT-qPCR_1.fwd | TCGTCCACAGCCAGGAAATC |
| hdeA_RT-qPCR_1.rev | CTGCCAGTTGTGAGCAATGC |
| hdeB_RT-qPCR_1.fwd | CATGCAACCGGGGTCATTG |
| hdeB_RT-qPCR_1.rev | GCTGTAGCGGCTTTGTCACT |
| yhiD_RT-qPCR_1.fwd | CACTGAACCATAAATACCCAGTTCG |
| yhiD_RT-qPCR_1.rev | TGTAGGTCTGACGACGGCAG |
| yhiD_RT-qPCR_2.fwd | GAAGCTGGATATCAATGGCGAC |
| yhiD_RT-qPCR_2.rev | CGGTATCAATGCTCGACTGG |
| dctR_RT-qPCR_1.fwd | TAAACGCTGCCTGGGGAAAG |
| dctR_RT-qPCR_1.rev | GTAACTGCCATTGATGAGTGCGAC |
| gade_RT-qPCR_1.fwd | ACGCTCAATATTTCGCAACAAAC |
| gade_RT-qPCR_1.rev | GTGATACCCAGGGTGACGATG |
| nrdF_RT-qPCR_1.fwd | ATCGCCTGACCAGCAATTTC |
| nrdF_RT-qPCR_1.rev | CGAGCAGCGTCAGGCCAGTA |
| nrdF_RT-qPCR_2.fwd | CGCAGAAATGGCGGAAGTGAATC |
| nrdF_RT-qPCR_2.rev | TCACATAAGAGGAGCCTGAACCG |
| proU_RT-qPCR_1.fwd | CGCTATCTTTGACAAAAAATATCAACTTTCTCG |
| proU_RT-qPCR_1.rev | GAATCTGAGGCAACCCCTGATGG |
| proV_RT-PCR_1.fwd | TCCGACGCCGAACTCCG |
| proV_RT-PCR_1.rev | ACGCAGTATTGTCCAGCACG |
| proV_RT-PCR_2.fwd | GGCGAATGATTATGTCCGTACCTTCTTC |
| proV_RT-PCR_2.rev | CGAAGCCAGGGGTTTTACGAAT |
| proW_RT-PCR_1.fwd | CCATTTCCGTCCCGTCTTCCAG |
| proW_RT-PCR_1.rev | GGCGATGAGAGCGAAAACGATAATC |
| proW_RT-PCR_2.fwd | GCCAGATGCTGTTCAAAGTTCAGTTAC |
| proW_RT-PCR_2.rev | GCGATGACCACCATAGAAAGGGC |
| proX_RT-PCR_1.fwd | GAAAGATCCGAAGATCGCCAAACTGTTC |
| proX_RT-PCR_1.rev | GTTGATCGCACCTTCGCAGC |
| proX_RT-PCR_2.fwd | TATCGTTGCCAACAAAGCCTGGG |
| proX_RT-PCR_2.rev | TAATGGCGTTCTGGGCGTTAATATCTG |
| ygaY_RT-qPCR_1.fwd | CTTTTCCCTTTCCGCCAGTTCG |
| ygaY_RT-qPCR_1.rev | CAGTAAGGTCATCGAGACAATCAGG |

Classified as Internal | Intern

| | |
|---|---|
| ygaY_RT-qPCR_2.fwd | GCCGCTCCACCTTTTAACTACAG |
| ygaY_RT-qPCR_2.rev | GGTGTGGTGCGATTTGCCCTTAT |
| Fw_hns1_RT_qPCR | GAACAACATCCGTACTCTTCGTGC |
| Re_hns1_RT_qPCR | CTTCTAATTTTTCCAGCATTTCTTCCAGC |
| Fw_stpA1_RT-qPCR | CGAATTCTCCATTGACGTTCTTGAAG |
| Re_stpA1_RT-qPCR | CTCTGCCAGTTCACGCTGC |
| rpoD_RT-qPCR.fwd | GACGAAGAAGATGGCGATGACGAC |
| rpoD_RT-qPCR.rev | TTCCTGAGCGGTAGCGTGACTG |
| rrsA_qRT-PCR_fwd | CTCTTGCCATCGGATGTGCCCA |
| rrsA_qRT-PCR_rev | CCAGTGTGGCTGGTCATCCTCTCA |
| dnaA_RT-qPCR_control_fwd | CGATCTAACGTACGTGAGCTGG |
| dnaA_RTqPCR_control_rev | GCACGAAGTCGATGGTGATCG |

Classified as Internal | Intern

#### Table S2. Primers for ChIP-qPCR

| name | internal name | Sequences (5'-3') |
|---|---|---|
| hdeD3 | Fw_hdeAB_yhiD15_ChIPqPCR | CATGGCTGATATTTTCCGTAGTCAGG |
| | hdeAB-yhiD15_MluCI | GATTACTTATTTAACCGCCCATAGC |
| hdeD1 | hdeD_RT-qPCR_1.fwd | CCGATGGTGTCTGTAACGC |
| | hdeD_RT-qPCR_1.rev | GCTTAACGAACAAACTGGCG |
| hdeD2 | hdeD_RT-qPCR_2.fwd | GCAATCCAGTTTATTGCCGTGC |
| | hdeD_RT-qPCR_2.rev | AATGCACCCACTACTGTGC |
| hdeAB1 | hdeAB-yhiD11_MluCI (Anchor primer1) | GAAGAAAATCCCCTGCTATCAATC |
| | Fw_hdeA_ChIP-qPCR1 | GAGTTACAGATGCATCAGATACTGC |
| hdeAB2 | Re_hdeA_ChIP-qPCR2 | CCATCAACATGACATATACAGAAAACCAGG |
| | Fw_hdeA_ChIP-qPCR2 | CCTCAACTATAAAGTGAAAGAGCCGTC |
| hdeA1 | hdeA_RT-qPCR_1.fwd | TCGTCCACAGCCAGGAAATC |
| | hdeA_RT-qPCR_1.rev | CTGCCAGTTGTGAGCAATGC |
| hdeB1 | hdeB_RT-qPCR_1.fwd | CATGCAACCGGGGTCATTG |
| | hdeB_RT-qPCR_1.rev | GCTGTAGCGGCTTTGTCACT |
| yhiD1 | yhiD_RT-qPCR_1.fwd | CACTGAACCATAAATACCCAGTTCG |
| | yhiD_RT-qPCR_1.rev | TGTAGGTCTGACGACGGCAG |
| yhiD2 | yhiD_RT-qPCR_2.fwd | GAAGCTGGATATCAATGGCGAC |
| | yhiD_RT-qPCR_2.rev | CGGTATCAATGCTCGACTGG |
| dctR1 | dctR_RT-qPCR_1.fwd | TAAACGCTGCCTGGGGAAAG |
| | dctR_RT-qPCR_1.rev | GTAACTGCCATTGATGAGTGCGAC |
| dctR2 | hdeAB-yhiD2_MluCI | CGTCGATCTCTTCTTCGTTC |
| | Fw_hdeAB_yhiD2_ChIPqPCR | CTTATAATTACCAGGGATACGATGTTC |
| nrdF1 | nrdF_RT-qPCR_1.fwd | ATCGCCTGACCAGCAATTTC |
| | nrdF_RT-qPCR_1.rev | CGAGCAGCGTCAGGCCAGTA |
| nrdF2 | nrdF_RT-qPCR_2.fwd | CGCAGAAATGGCGGAAGTGAATC |
| | nrdF_RT-qPCR_2.rev | TCACATAAGAGGAGCCTGAACCG |
| proU1 | proU_RT-qPCR_1.fwd | CGCTATCTTTGACAAAAAATATCAACTTTCTCG |
| | proU_RT-qPCR_1.rev | GAATCTGAGGCAACCCCTGATGG |
| proV3 | proV_RT-qPCR_3.fwd | GGTAATATATCGACATAGACAAATAAAGGAATCTTTCTATTGCATG |
| | proV_RT-qPCR_3.rev | CCAGAATTTGTTCTTTTGAAAGTCCTTGTTCG |
| proV1 | proV_RT-PCR_1.fwd | TCCGACGCCGAACTCCG |
| | proV_RT-PCR_1.rev | ACGCAGTATTGTCCAGCACG |
| proV2 | proV_RT-PCR_2.fwd | GGCGAATGATTATGTCCGTACCTTCTTC |
| | proV_RT-PCR_2.rev | CGAAGCCAGGGGTTTTACGAAT |
| proW1 | proW_RT-PCR_1.fwd | CCATTTCCGTCCCGTCTTCCAG |
| | proW_RT-PCR_1.rev | GGCGATGAGAGCGAAAACGATAATC |
| proW2 | proW_RT-PCR_2.fwd | GCCAGATGCTGTTCAAAGTTCAGTTAC |
| | proW_RT-PCR_2.rev | GCGATGACCACCATAGAAAGGGC |

Classified as Internal | Intern

| | | |
|---|---|---|
| proX1 | proX_RT-PCR_1.fwd | GAAAGATCCGAAGATCGCCAAACTGTTC |
| | proX_RT-PCR_1.rev | GTTGATCGCACCTTCGCAGC |
| proX2 | proX_RT-PCR_2.fwd | TATCGTTGCCAACAAAGCCTGGG |
| | proX_RT-PCR_2.rev | TAATGGCGTTCTGGGCGTTAATATCTG |
| ygaY1 | ygaY_RT-qPCR_1.fwd | CTTTTCCCTTTCCGCCAGTTCG |
| | ygaY_RT-qPCR_1.rev | CAGTAAGGTCATCGAGACAATCAGG |
| ygaY2 | ygaY_RT-qPCR_2.fwd | GCCGCTCCACCTTTTAACTACAG |
| | ygaY_RT-qPCR_2.rev | GGTGTGGTGCGATTTGCCCTTAT |
| dnaA | dnaA_RT-qPCR_control_fwd | CGATCTAACGTACGTGAGCTGG |
| | dnaA_RTqPCR_control_rev | GCACGAAGTCGATGGTGATCG |
| appB1 | FW_appB_ChIPqPCR_NC1 | CTGGGAAGGCAACCAGGTC |
| | RE_appB_ChIPqPCR_NC1 | GGAAAACGCCGCTGCATAC |

#### Table S3. Primers for *hdeAB* operon standard library preparation

| Fragment No. | Internal name | Sequences (5'-3') |
|---|---|---|
| 1 | Fw_IC1_Probe2_1 | GTATCCGTTTTGCCATTGATAACGTTGATAACCTTCCCACCAAAGCGCGCTTGTTGACCAACATATAACCCC |
| 1 | Re_IC1_Probe2_1 | CAAAGGCAATAACCAACCTGATATTCAAAAAAGTTTTGTTGCTGTTCATAACCAGCCGGGGTTATATGTTGGTC |
| 1 | Fw_IC1_Probe2_2 | GGTTATTGCCTTTGATATTTTGCGGATTTCGACACTGAGGTTATAACCTGGTTTTCTGTATATGTCATG |
| 2 | Fw_IC2_Probe2_1 | CGTCGATCTCTTCTTCGTTCTGTATATGAACGACATTACCTTTACTCAGAATG |
| 2 | Re_IC2_Probe2_1 | GAAATACCAGGGATACGATGTTCTTCACCGCGATGAAAAACATTCTGAGTAAAGGTAATG |
| 2 | Fw_IC2_Probe2_2 | CATCGTATCCCTGGTATTTCGACACTGAGGTTATAACCTGGTTTTCTGTATATGTCATG |
| 3 | Fw_IC3_Probe2_1 | GTAACTGCCATTGATGAGTGCGACAAAATCAATGATAGTGATAGTCCGCGGAACAAAGGTCACCGGCACTTTC |
| 3 | Re_IC3_Probe2_1 | TAAAACCTGTCCATGTCATTATTTTCTCCCCCTTTAATATTAAACGCTGCCTGGGGAAAGTGCCGGTGAC |
| 3 | Fw_IC3_Probe2_2 | GACATGGACAGGTTTTAATCGTTCAATTTCGACACTGAGGTTATAACCTGGTTTTCTGTATATGTCATG |
| 4 | Fw_IC4_Probe2_1 | CTGATTTAGTTTCGTTGCAGGAAAATGAAGATCATGAAGTGGTCGCCATTGATATCCAGCTTCATGCCAC |
| 4 | Re_IC4_Probe2_1 | CACCCCTGCCATACCTTTTAATAATCGCAACAGGTCTTCTATCGACGTTGTGGCATGAAGCTGGATATC |
| 4 | Fw_IC4_Probe2_2 | GGTATGGCAGGGGTGAAAGGCGTTTCAATTTCGACACTGAGGTTATAACCTGGTTTTCTGTATATGTCATG |
| 5 | Fw_IC5_Probe2_1 | GTAGGTCTGACGACGGCAGCGGATATCTGGGTGACCGCCGCCATAGGTATGGTTATTGGCAGCGGTATGTAC |
| 5 | Re_IC5_Probe2_1 | CTTCCAGCACCAACAAGGTCATCACTGAACCATAAATACCCAGTTCGTACATACCGCTGCCAATAAC |
| 5 | Fw_IC5_Probe2_2 | CTTGTTGGTGCTGGAAGTCTTCCATCAATTTCGACACTGAGGTTATAACCTGGTTTTCTGTATATGTCATG |
| 6 | Fw_IC6_Probe2_1 | GAGTAGCAAGTTGAGCCATCTTGCTGCTCCTTTTTGCATTTTTATATGACAGC |
| 6 | Re_IC6_Probe2_1 | CCAGGTTATAACCTCAGTGTCGAAATTCTGCTGTCATATAAAAATGC |
| 6 | Fw_IC6_Probe2_2 | GAATTTCGACACTGAGGTTATAACCTGGTTTTCTGTATATGTCATG |
| 7 | Fw_IC7_Probe2_1 | GGTGGATGCTGCATGAAGAAACAGTATATAAAGGTGGCGATACCGTTACTTTAAATG |
| 7 | Re_IC7_Probe2_1 | GTTATAACCTCAGTGTCGAAATTTGAGTGAGATCGGTTTCATTTAAAGTAACGGTATCGCC |
| 7 | Fw_IC7_Probe2_2 | AATTTCGACACTGAGGTTATAACCTGGTTTTCTGTATATGTCATGTTG |
| 8 | Fw_IC8_Probe2_1 | CATCTCTCCGTAAAGCGTTTATTTTTATGGGCGCTGTAGCGGCTTTGTCACTGGTGAACG |
| 8 | Re_IC8_Probe2_1 | GTCATATCTTTAGCGGATTCATTGGCTGCCAACGCAGATTGTGCGTTCACCAGTGACAAAGCCG |
| 8 | Fw_IC8_Probe2_2 | CCGCTAAAGATATGACCTGCCAGGAATTTCGACACTGAGGTTATAACCTGGTTTTCTGTATATGTCATG |
| 9 | Fw_IC9_Probe2_1 | GTAACCCCAGCTATCGTTCAGGCTTGTACTCAGGATAAACAAGCCAACTTTAAAG |

Classified as Internal | Intern

| | | |
|---|---|---|
| 9 | Re_IC9_Probe2_1 | GAAATTTTGTCCCATTCGCCTTTAACTTTATCTTTAAAGTTGGCTTGTTTATC |
| 9 | Fw_IC9_Probe2_2 | GGCGAATGGGACAAAATTTCGACACTGAGGTTATAACCTGGTTTTCTGTATATGTCATG |
| 11 | Fw_IC11_Probe2_1 | GAAGAAAATCCCCTGCTATCAATCTATGCCAAAAACGCGTCTAAG |
| 11 | Re_IC11_Probe2_1 | CCTCAGTGTCGAAATTTTTATTAAATCGACTGCATTCTTAGACGCGTTTTTGG |
| 11 | Fw_IC11_Probe2_2 | AAAATTTCGACACTGAGGTTATAACCTGGTTTTCTGTATATGTCATG |
| 12 | Fw_IC12_Probe2_1 | GATACACAGCAACCCGACGATAAACAGCAGCACGGCAATAAACTGGATTGCTCTGC |
| 12 | Re_IC12_Probe2_1 | GAAGTTTGATCTGGAGATGCTTAAAAAACATCGCAGAGCAATCCAGTTTATTG |
| 12 | Fw_IC12_Probe2_2 | CATCTCCAGATCAAACTTCAAAATTTCGACACTGAGGTTATAACCTGGTTTTCTGTATATGTCATG |
| 13 | Fw_IC13_Probe2_1 | CAAAAATGCCCAGCTCCGGTGCGCGGATGAAGAAATAGCCGATCAATAAATAG |
| 13 | Re_IC13_Probe2_1 | CTGGCCGGTATTATCCGGTTTCCTCGTCGCAGTCGCCTATTTATTGATCGGC |
| 13 | Fw_IC13_Probe2_2 | GAAACCGGATAATACCGGCCAGAAATTTCGACACTGAGGTTATAACCTGGTTTTCTGTATATGTCATG |
| 14 | Fw_IC14_Probe2_1 | GTAGCGGAGAGAGTATTAATCGGATCATAGTCACATCAAGTGAC |
| 14 | Re_IC14_Probe2_1 | GAAATTACCCCGGTTGTCACCCGGATCATAGTCACTTGATGTGACTATG |
| 14 | Fw_IC14_Probe2_2 | GACAACCGGGGTAATTTCGACACTGAGGTTATAACCTGGTTTTCTGTATATGTCATG |

Classified as Internal | Intern

#### Table S4. Primers for 3C-qPCR

| Fragment No. | Internal name | Sequences (5'-3') |
|---|---|---|
| 1 | hdeAB-yhiD1_MluCI | GTATCCGTTTTGCCATTGATAAC |
| 2 | hdeAB-yhiD2_MluCI | CGTCGATCTCTTCTTCGTTC |
| 3 | hdeAB-yhiD3_MluCI | GTAACTGCCATTGATGAGTGC |
| 4 | hdeAB-yhiD4_MluCI | CTGATTTAGTTTCGTTGCAGG |
| 5 | hdeAB-yhiD5_MluCI | GTAGGTCTGACGACGG |
| 6 | hdeAB-yhiD6_MluCI | GAGTAGCAAGTTGAGCCATC |
| 7 | hdeAB-yhiD7_MluCI | GGTGGATGCTGCATGAAG |
| 8 | hdeAB-yhiD8_MluCI | CATCTCTCCGTAAAGCG |
| 9 | hdeAB-yhiD9_MluCI | GTAACCCCAGCTATCGTTCAG |
| 10 | hdeAB-yhiD10_MluCI(Anchor primer) | GCATCTGTAACTCATTGTATTGAAA |
| 11 | hdeAB-yhiD11_MluCI | GAAGAAAATCCCCTGCTATCAATC |
| 12 | hdeAB-yhiD12_MluCI | GATACACAGCAACCCGAC |
| 13 | hdeAB-yhiD13_MluCI | CAAAAATGCCCAGCTCCG |
| 14 | hdeAB-yhiD14_MluCI | GTAGCGGAGAGAGTATTAATCG |

#### Table S5. Details of TaqMan-probe

| Name | Sequences (5'-3') | Purification |
|---|---|---|
| TaqMan probe_*hdeAB* | 6-FAM/TGAGGTTATAACCTGGTTTTCT/ZEN/GTATATGTCATGTTGATGG/IBFQ | RP-HPLC |
| TaqMan probe_*proVWX* | 6-FAM/CATGCAATA/ZEN/GAAAGATTCCTTTATTTGTCTATGTCGAT/IBFQ | RP-HPLC |

Classified as Internal | Intern
